# Bacteriocin Diversity and Antiviral Potential of *Lactiplantibacillus pentosus from* Fermented Rice

**DOI:** 10.64898/2026.04.22.720035

**Authors:** Athira Cheruvari, Rajagopal Kammara

**Affiliations:** Department of Biochemistry, CSIR-Central Food Technological Research Institute, Mysore, 570020, India; Academy of Scientific and Innovative Research (AcSIR), Ghaziabad-201002, India

**Keywords:** Comparative genomics, *Lactobacillus pentosus*, KEGG pathway, antimicrobials, Whole Genome Sequencing

## Abstract

Fermentation is a long-standing tradition on the Indian subcontinent, shaped by local cultures and resources. Regional ingredients create a variety of fermented foods rich in probiotics. Lactiplantibacillus pentosus, found in foods like olives, grains, and vegetables, is a key strain. We studied the genome of *L. pentosus* from traditional fermented rice in Himachal Pradesh, India, and compared it to 95 global genomes. Our focus was on antimicrobial compounds, especially bacteriocins, which improve food safety. KEGG analysis revealed important metabolic proteins, while BPGA showed genome sizes of 3.4-4 Mb and GC content of 45.5-46%. The Indian L. pentosus strain (krglsrbmofpi2) uniquely contains three bacteriocins: Pentoplantaricin-EF, Pentobovicin, and Pentopediocin. Docking studies assessed their antiviral potential. Pentoplantaricin-EF showed the strongest binding to the prefusion 2019-nCoV spike glycoprotein (PDB 6VSB), while Pentobovicin had the highest affinity for the Omicron spike protein (PDB 7T9J) and the Hepatitis E virus E2 domain (PDB 3RKC).

## INTRODUCTION

Traditional fermented foods represent one of the oldest and most cost-effective methods of food preservation, valued for their nutritional richness and probiotic content, including lactic acid bacteria (LAB) and bifidobacteria [1]. In Indian cuisine, these foods are categorized by source — cereals, dairy, vegetables, legumes, meat, and bamboo shoots — with quality dependent on microbial composition of raw materials [2]. Their global popularity has grown alongside research linking consumption to numerous health benefits [3]. LAB play a central role in fermentation by producing lactic acid while enhancing flavor, aroma, and nutritional quality [4,5], and are associated with pathogen inhibition and immune modulation, though their mechanisms remain incompletely understood [6]. Probiotic strains must survive harsh gastrointestinal conditions to be effective [7], and research has expanded their applications to nanotechnology, cancer treatment, and functional foods [8,9]. LAB also produces bacteriocins — antimicrobial peptides that inhibit pathogens and spoilage bacteria [10]. Though sensitive to gastric enzymes, bacteriocins are generally safe and are being investigated as antibacterial agents [11]; Lactobacillus-derived examples such as nisin are already commercially available [12]. This manuscript examines Lactobacillus genomes to characterize bacteriocin diversity, genomic location, biosynthesis, transport, self-immunity, and target specificity, highlighting their potential as natural antibiotics for germ-free food production.

## METHODS

### Bacterial strain and culture condition

*Lactiplantibacillus pentosus* was isolated from Himachal Pradesh fermented rice and subsequently preserved as frozen stocks at a temperature of −80°C in MRS broth supplemented with 50% (v/v) glycerol. The strain was identified by 16S rRNA sequencing and Whole genome sequencing. The strain was grown in MRS Broth (at 37°C, 24 hrs) in aerobic conditions [13].

### Genome Sequencing

Genomic DNA was isolated using the GeneJet Bacterial Genomic DNA Purification Kit from Thermo Fisher Scientific. The sequencing was performed using Illumina HiSeq 1000 technology by Sandor Speciality Diagnostics in Hyderabad. The resulting data was submitted to the NCBI (National Centre for Biotechnology Information) [14,15].

### Functional annotations

The proteins encoded in the genome of the *L. pentosus* Indian strain, krglsrbmofpi2, were annotated and categorized using Blast KOALA (BLAST-based KO annotation and KEGG mapping) against the KEGG Kyoto Encyclopaedia of Genes and Genomes (KEGG) database [16]. This database displays proteins that are involved in various metabolic processes [17].

### Genome Data collection and classification

As of December 2023, 98 *Lactiplantibacillus pentosus* genomes were available in NCBI, of which 96 — including an isolate from Himachal Pradesh, India — had comprehensive genome and protein sequence documentation. These genomes were downloaded, categorized by size, GC content, origin, and source, and 95 were compared with the Indian isolate. A global distribution map was created using MapChart software based on isolation or submission sites [18] (accessed 30 December 2023, https://www.mapchart.net/), and a graph was generated showing annual *L. pentosus* genome releases through December 2023.

### Comparative Bacterial Pan-Genome Analysis

Pan- and core-genome analysis of 96 *L. pentosus* strains was performed using BPGA [19] to identify strain-specific features and genomic diversity. Protein files from all 96 genomes were downloaded from NCBI and used as input; BPGA clustered genes into families via USEARCH at a 50% sequence-similarity cutoff to define the pan-genome. COG categories and KEGG pathway distributions were assessed through BPGA’s functional analysis module, and evolutionary analysis was based on concatenated core gene alignment using a binary pan-matrix depicting gene presence or absence across genomes [20, 21].

### Prediction of Genes Related to Antibacterial Activity

The BAGEL-4 online tool was used to identify bacteriocin genes within 96 genomes of *L. pentosus*. This web server is designed for the identification and visualization of gene clusters in prokaryotic DNA associated with bacteriocin biosynthesis [22, 23]. The predicted bacteriocins were then categorized according to their types, and histograms were generated to illustrate their distribution across the genomes. Additionally, the bacteriocin sequences among the genomes were analyzed to identify similarities and differences.

### Molecular Docking studies

We investigated the potential antiviral activity of three novel bacteriocins—Pentobovicin (Bovicin), Pentopediocin (Pediocin, Pd), and Pentoplantaricin_EF EF (Plantaricin EF, Pln) produced by *Lactiplantibacillus pentosus* (krglsrbmofpi2), a strain isolated from fermented rice in Himachal Pradesh. Using *in silico* approaches, we analyzed their interactions with three viral targets: the Hepatitis E Virus Capsid Protein E2s Domain, the Cryo-EM structure of the SARS-CoV-2 Omicron spike protein, and the Prefusion 2019-nCoV spike glycoprotein with a single receptor-binding domain. This study aims to investigate the potential of these novel bacteriocins as antiviral agents, providing insight into their possible mechanisms of action and broadening their therapeutic applications.

### Ligand preparation

A bacterial strain isolated from fermented rice in Himachal Pradesh, India, was identified and confirmed as *Lactiplantibacillus pentosus* (krglsrbmofpi2), through genomic characterization. The presence and sequences of its bacteriocins—Bovicin/Putative bacteriocin (Pentobovicin), Pediocin (Pentopediocin), and Plantaricin_EF (Pentoplantaricin EF) were detected using BAGEL 4, a specialized tool for identifying bacteriocin-encoding genes [24, 25]. The primary amino acid sequences of the identified bacteriocins were used to predict three-dimensional structures using I-TASSER (Iterative Threading Assembly Refinement; accessed June 3–17, 2024), which employs threading alignments and structural assembly simulations based on known protein templates [26, 27, 28]. Model confidence was assessed via C-score, and the most reliable structures were selected as ligands for molecular docking studies to explore their potential antiviral mechanisms.

### Target selection

Target selection focused on viral proteins critical for host recognition, infection, and immune evasion, driven by the rising threat of COVID-19 and Hepatitis and the need for novel antiviral agents. The 3D structures of three key proteins were retrieved from the Protein Data Bank (PDB) [29]: the prefusion 2019-nCoV spike glycoprotein (PDB ID: 6VSB) [30, 31], the SARS-CoV-2 Omicron spike protein (PDB ID: 7T9J) [32, 33, 34], and the Hepatitis E Virus Capsid Protein E2s Domain (PDB ID: 3RKC) [35]— all central to viral attachment, entry, and immune evasion, making them ideal for *in silico* screening.

### Molecular Docking

Molecular docking simulations were performed using ClusPro, which applies FFT-based rigid-body docking. The study evaluated binding affinities and compatibility of *L. pentosus* krglsrbmofpi2-derived bacteriocins—Pentobovicin, Pentopediocin, and Pentoplantaricin_EF—with viral proteins: Prefusion 2019-nCoV spike glycoprotein (6VSB), SARS-CoV-2 Omicron spike protein (7T9J), and Hepatitis E virus capsid protein E2S domain (3RKC). ClusPro default parameters were applied, involving FFT-based rigid-body docking, energy-based filtering of low-energy conformations, clustering of docked poses, and ranking by balanced, electrostatic, hydrophobic, and van der Waals interactions. Top-ranked models, selected by cluster size and binding energy, were used for further interaction analysis [36, 37, 38, 39, 40].

### Analysis of Protein-Protein Interactions

The resulting docked complexes were further analyzed to evaluate the nature of interactions at the protein-protein interface. Key interaction parameters, including hydrogen bonding, hydrophobic contacts, and electrostatic interactions, were examined to determine the binding stability and specificity of the bacteriocins to the viral proteins. PyMOL 3.1 software was used to visualize the docked complexes and assess the molecular interactions between the recombinant bacteriocins and the viral targets. Structural compatibility, binding affinities, and interaction sites were analyzed to infer possible inhibitory mechanisms by which these bacteriocins could disrupt viral functionality [41, 42, 43].

## Results

*Lactobacillus pentosus* strain krglsrbmofpi2 (Cheruvari and Rajagopal 2024) was isolated from fermented rice in Himachal Pradesh, India. Its genome contains 3,418 genes (3,192 coding), deposited in NCBI under accession GCA_009295675.1, and annotated using PGAP. Functional annotation with the BlastKOALA tool (KEGG Orthology database) successfully annotated 51.3% of genes across metabolic and other pathways (Suppl. Figure 1, Suppl. Table 1). Of 98 available genomes (December 2023), 96 with complete details were used. Genome sizes range from 3.4 to 4 Mb with GC content between 45.5% and 46%; larger genomes confer environmental adaptability, while higher GC content may increase reproductive energy costs [44]. A world map of strain distribution was generated (Figure 1a), with strains classified by origin — plant, animal, human, or other (Suppl. Tables 2 and 3); *L. pentosus* is predominantly of plant origin. Annual genome releases through December 2023 are shown in Suppl. Figures 2 and 3, with 2021 recording the highest number of releases. BPGA analysis of the 96 genomes identified core and accessory genes (Suppl. Table 4, Suppl. Fig. 3a, b), with KEGG analysis revealing carbohydrate metabolism as the dominant pathway (Suppl. Fig. 3c), and COG distributions for core, accessory, and unique genes shown in Suppl. Fig. 3d, e. Phylogenetic analysis (Suppl. Figure 1b) confirmed the Indian isolate as newly reported, with distinct bacteriocin gene arrangements. Genome mining identified six bacteriocin types: pediocin, plantaracin, bovicin, sactipeptides, LAPs, and pentocin (Suppl. Fig. 4a–c, Suppl. Table 5). Pediocin was most prevalent (91 strains), followed by plantaracin (62), sactipeptides (59), bovicin (14), and LAPs (4) (Suppl. Fig. 4). Among plantaracins, four subtypes were identified: A (53 strains), NC8 alpha-beta (22), EF (9), and S-alpha-beta (4) (Suppl. Fig. 4B). Strains produce one to five bacteriocins: 31 strains produce 2–3, ten produce one, eight produce four, and six produce five (Suppl. Fig. 4C). The Indian strain (krglsrbmofpi2) uniquely produces the combination Pd, PlnF, and Bov — distinct from all other observed combinations (Suppl. Fig. 4d). We conducted a thorough analysis of the *L. pentosus* strain genome, followed by sequence alignment of Plantaracin E and F. Our investigation revealed a significant mutation in which valine (indicated in red) replaced cysteine, highlighting the potential impact of this change. No mutations were observed in plantaracin F. Overall, the bacteriocin combination and sequence are unique. Supplementary Figure 5a, b shows the sequence of pediocin and bovicin; found similarity but no variation. Supplementary Figure 5 d, e represents the bacteriocin gene plnEF, Pediocin, and bovicin arrangement and their orientation.

**Figure 1.**
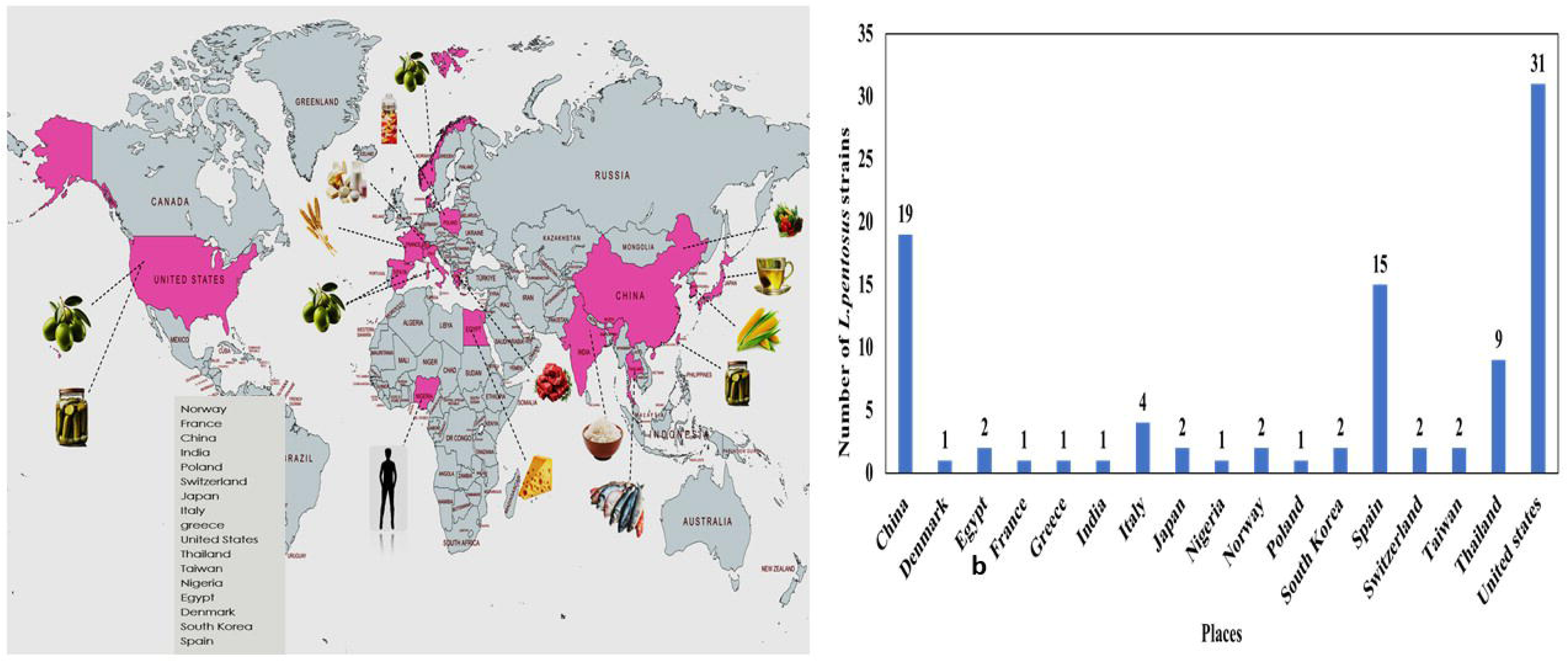
a, b: Depicts worldwide distribution of *L. pentosus* strain and its major sources (Data from NCBI up to December 2023).

### Molecular Docking studies Identification of Novel Bacteriocins

Genomic analysis of *Lactiplantibacillus pentosus* (krglsrbmofpi2) identified three bacteriocins — Bovicin/Putative bacteriocin, Pediocin, and Plantaricin_EF — using BAGEL 4 (Cheruvari and Kammara 2024). Sequence comparisons confirmed that while these share similarity with known bacteriocins, they are not 100% identical to any characterized variant. Since even a single amino acid difference can significantly affect activity, they were designated as novel variants — Pentobovicin, Pentopediocin, and Pentoplantaricin_EF — reflecting their *L. pentosus* origin and aligning with current classification guidelines [45].

### Structure Prediction and Validation

3D structures of Pentobovicin, Pentopediocin, and Pentoplantaricin_EF were predicted using I-TASSER and evaluated by C-score (range: -5 to 2; higher = more reliable) and TM-score (range: 0–1; >0.5 = reliable fold). Secondary structure (α-helices, β-strands, coils), solvent accessibility, and normalized B-factor analyses provided insights into folding, stability, and function (suppl. Fig. 6a–c) [46].

I-TASSER generated five models per bacteriocin; the highest C-score model was selected. Pentopediocin had the best predicted structure (C-score: 0.59, TM-score: 0.64 ± 0.14), with the top PDB hit provided in Suppl. Table 6. Suppl. Figure 6 illustrates predicted secondary structures, normalized B-factors, and solvent accessibility for all three bacteriocins. C-score (range: -5 to 2; higher = more reliable), TM-score (>0.5 = correct fold), and PDB Hit (closest PDB match used for prediction) are detailed in Suppl. Table 6.

High-confidence models of Pentobovicin, Pentoplantaricin_EF, and Pentopediocin were selected as ligands and visualized in PyMOL (v3.0; Suppl. Figs. 7a–f). Viral target structures — prefusion 2019-nCoV spike glycoprotein (PDB: 6VSB), SARS-CoV-2 Omicron spike protein (PDB: 7T9J), and Hepatitis E Virus Capsid Protein E2S Domain (PDB: 3RKC) — were retrieved from PDB and visualized in PyMOL (Suppl. Fig. 8a–d).

### Molecular Docking

ClusPro docking yielded 10 clustered poses per interaction; top-ranked clusters (most negative energy = most favorable) were selected for analysis, with binding energies summarized in Suppl. Table 7.

Pentobovicin binding energies: -971.7 kcal/mol with prefusion 2019-nCoV spike glycoprotein (PDB: 6VSB; Fig. 2A), -1153.4 kcal/mol with SARS-CoV-2 Omicron spike protein (PDB: 7T9J; Fig. 2B), and -928.8 kcal/mol with Hepatitis E Virus Capsid Protein E2S domain (PDB: 3RKC; Fig. 2C).

**Figure 2:**
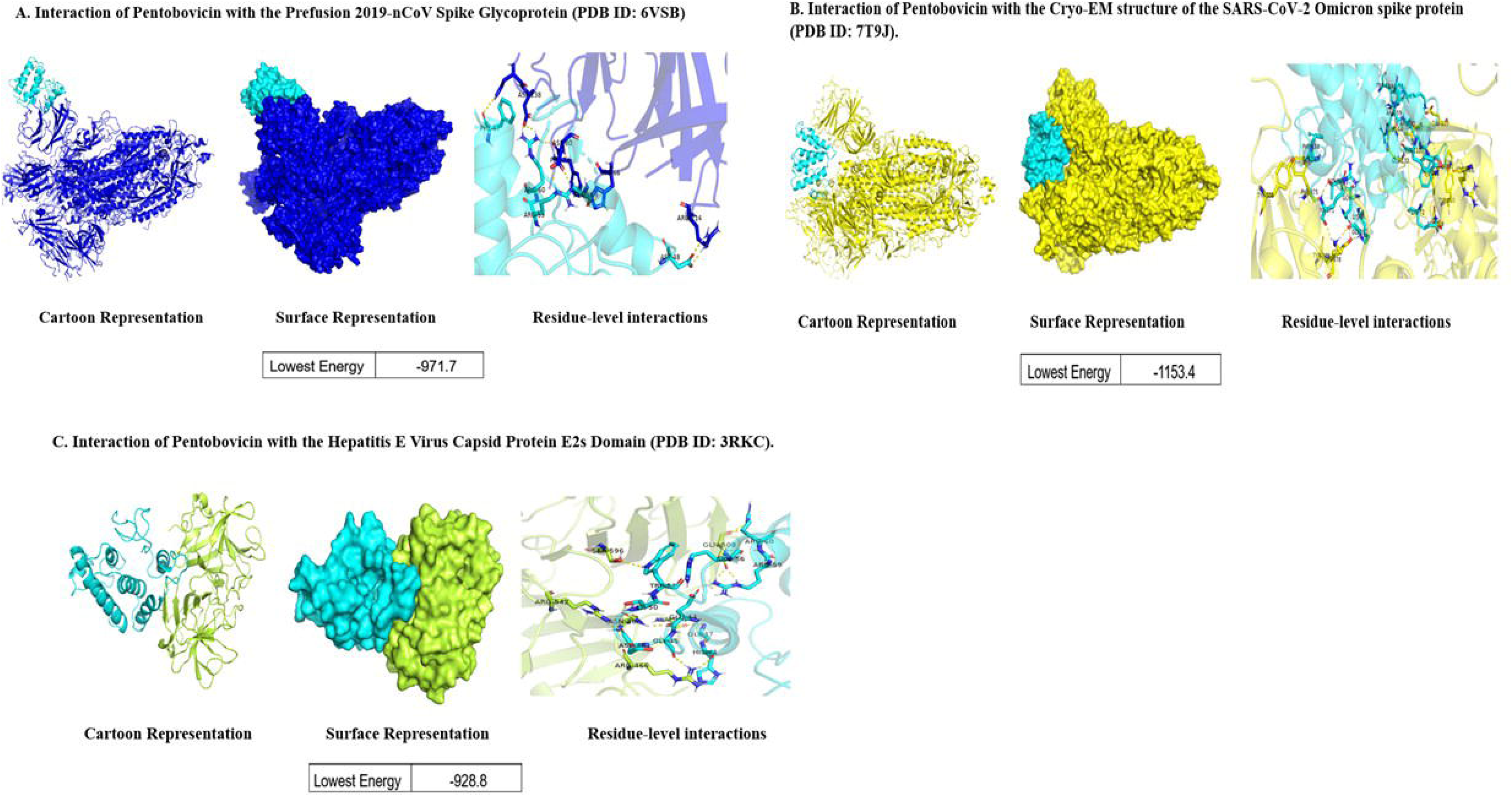
Representing interaction of Pentobovicin with different viral proteins. A) Interaction of Pentobovicin with the Prefusion 2019-nCov Spike glycoprotein (PDB ID:6VSB), B) Interaction of Pentobovicin with cryo-EM structure of the SARS-Cov-2 Omicron spike protein (PDB ID: 7T9J), C) Interaction of Pentobovicin with the Hepatitis E Virus Capsid Protein E2s domain (PDB ID: 3RKC).

Pentoplantaricin_EF showed the highest overall binding affinity, with -1439.5 kcal/mol against the prefusion 2019-nCoV spike glycoprotein (PDB: 6VSB; Fig. 3A), suggesting strong interaction with this target and potential influence on its structural stability. For PDB: 7T9J, Pentobovicin ranked highest, followed by Pentoplantaricin_EF and Pentopediocin (Fig. 3A–C); the same order held for PDB: 3RKC. Pentopediocin binding energies: -887.8 kcal/mol (PDB: 6VSB; Fig. 4a), -975.2 kcal/mol (PDB: 7T9J; Fig. 4b), and -869.2 kcal/mol (PDB: 3RKC; Fig. 4c; Suppl. Table 7). Variations in binding affinities reflect differences in molecular recognition that may influence antiviral efficacy (Figs. 2–4).

**Figure 3:**
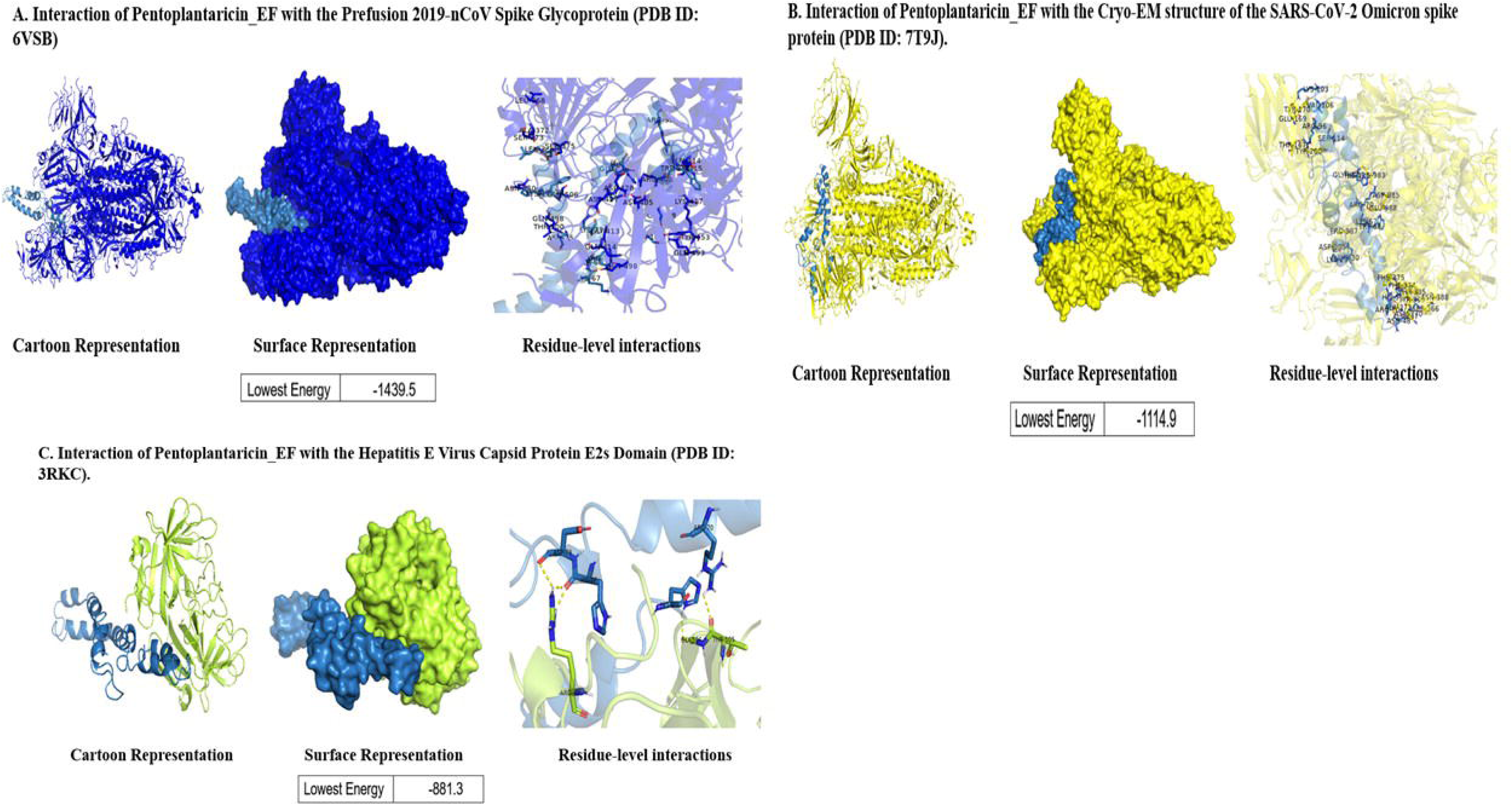
Representing interaction of Pentoplantaracin-EF with different viral proteins. A) Interaction of Pentoplantaracin-EF with the Prefusion 2019-nCov Spike glycoprotein (PDB ID:6VSB), B) Interaction of Pentoplantaracin-EF with cryo-EM structure of the SARS-Cov-2 Omicron spike protein (PDB ID: 7T9J), C) Interaction of Pentoplantaracin-EF with the Hepatitis E Virus Capsid Protein E2s domain (PDB ID: 3RKC).

**Figure 4:**
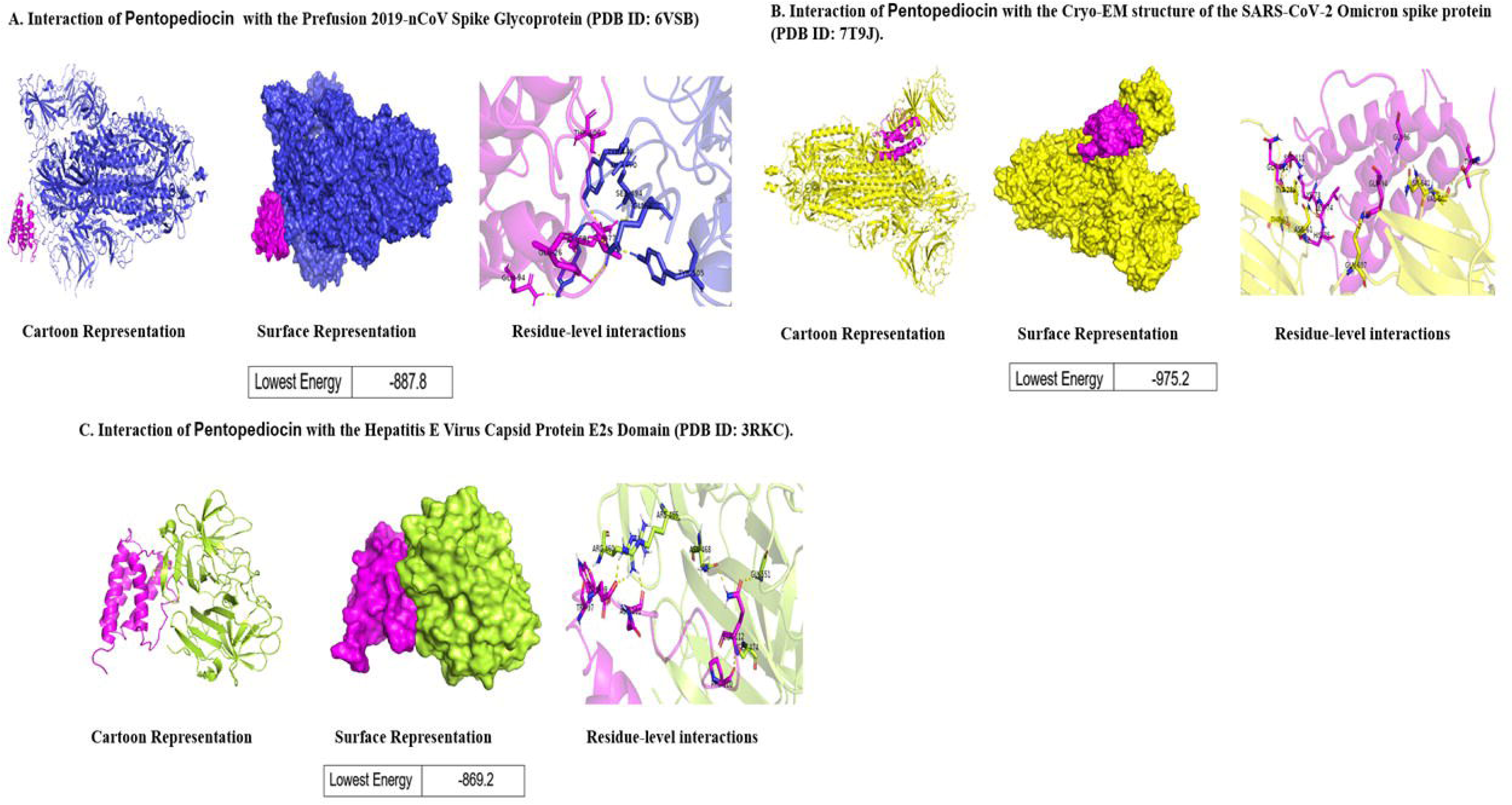
Representing interaction of Pentopediocin with different viral proteins. A). Interaction of Pentopediocin with the Prefusion 2019-nCov Spike glycoprotein (PDB ID:6VSB), B) Interaction of Pentopediocin with cryo-EM structure of the SARS-Cov-2 Omicron spike protein (PDB ID: 7T9J), C) Interaction of Pentopediocin with the Hepatitis E Virus Capsid Protein E2s domain (PDB ID: 3RKC).

## Discussion

Comparative genomics of *L. pentosus* reveals substantial diversity in bacteriocin production, shaped by geography and culture [47, 48, 49, 50]. Li et al. (2017) [51] identified genes underlying Weissellicin 110 production in *W. cibaria* 110, and Li et al. (2021) [52] found stress resistance and bacteriocin production genes in *P. acidilactici*, with implications for food preservation and probiotics.

These genomic comparisons underscore that regional differences in fermentation substrates shape distinct microbial populations and metabolic capabilities. Specifically, geography and culture drive unique functional genomic arrangements, suggesting novel antimicrobial properties. Regional fermentation substrates drive distinct microbial traits, as shown by Wang et al. (2020) [53] for *L. ruminis* GIT adaptation and Evanovich et al. (2019) [54] for *L. plantarum* pan-genome structure and environmental adaptability. These findings align with [55] Garcia-Gonzalez et al. (2022), who used comparative genomics to identify probiotic markers, bacteriocin production genes, and stress tolerance traits in *L. plantarum* strains from the human gut and fermented foods [56].

LAB bacteriocins inhibit drug-resistant Staphylococcus in goat milk cheeses [57]. BPGA analysis of 96 *L. pentosus* strains confirmed wide genetic diversity, reflecting adaptability and metabolic versatility in fermented food environments. Genomic diversity in Lactobacillus spp. underpins their fermentation and probiotic functions [58], providing a foundation for engineering *L. pentosus* strains with enhanced antimicrobial properties. Furthermore*, L. plantarum* produces various bacteriocins, notably NC8, which exhibit strong activity against Staphylococcus spp. and synergize with antibiotics. Recent studies also show the antiviral potential of PLNC8 αβ against flaviviruses and coronaviruses. Pentoplantaricin_EF showed the highest binding affinity (-1439.5 kcal/mol) against the prefusion 2019-nCoV spike glycoprotein (PDB: 6VSB; Fig. 11), consistent with antimicrobial peptides interfering with viral attachment [59, 60, 61, 62].

Against the SARS-CoV-2 Omicron spike protein (PDB: 7T9J), Pentobovicin showed the strongest binding (-1153.4 kcal/mol), followed by Pentopediocin (-975.2 kcal/mol) — comparable to or more potent than heterocyclic compounds reported by Akman et al. (2023). Against the Hepatitis E virus capsid protein E2S domain (PDB: 3RKC), Pentobovicin showed the strongest interaction. Pentoplantaricin_EF and Pentopediocin followed, reflecting differential molecular recognition. Similar trends have been reported for natural biomolecules against Dengue, Ebola, Zika, and SARS-CoV-2 [63, 64] demonstrated strong docking of bacteriocin- and defensin-derived peptides with HEV capsid proteins. This study highlights *L. pentosus* genetic diversity and the role of regional strains in producing unique antimicrobial compounds, with novel bacteriocins from the Indian strain offering opportunities for food safety and health applications. This study identified three novel bacteriocins — Pentobovicin, Pentopediocin, and Pentoplantaricin_EF — from *L. pentosus*, with docking analyses indicating antiviral potential against SARS-CoV-2 and Hepatitis E, supporting bacteriocins as alternatives to standard antivirals.

## Supporting information

statistical data

## Acknowledgements

We acknowledge CSIR-UGC for providing a fellowship to Athira Cheruvari, and the director for providing the resources.

We thank the Director CSIR-CFTRI, and AcSIR for their constant support during the project’s development.

## Data availability / Nucleotide sequence accession numbers

*L. pentosus* krglsrbmofpi2’s draft genome sequencing data has been released to GenBank with the accession number GCA_009295675.1 (Bio-Project accession number PRJNA576968 and BioSample accession number SAMN13015363). The isolated strain’s 16S rRNA gene sequence has the NCBI accession number MN165450 (it had previously been classified under *Lactobacillus plantarum*). Initially submitted as *Lactobacillus pentosus*, the genome eventually became *Lactiplantibacillus pentosus*. The strain has been identified as krglsrbmofpi2.

## CRediT authorship contribution statement

**KRG and AC:** Writing – original draft, Methodology, Investigation, Formal analysis, Data curation. **Xiao Ma:** Resources, Formal analysis, Data curation. **AC:** Visualization, Methodology. **KRG and AC:** Writing – review & editing, Validation, Software. **AC:** Resources, Data curation. **KRG:** Resources, Data curation. **KRG:** Writing – review & editing, Visualization, Investigation, Formal analysis, Conceptualization.

## Contributions

Dr KR designed the study. AC performed the experiments and collected the data. KR and AC analyzed the data. AC and KR wrote the original draft of the manuscript. AC, and KR reviewed and edited the manuscript. All authors have read and agreed to the final version of the manuscript.

## Corresponding author

Correspondence to K. Rajagopal

## Ethics declarations

### Ethical approval and consent to participate

Ethical approval was not required for this study as it did not involve vertebrate animals.

### Consent for publication

All authors consent to this publication.

## Supplementary Table Legends

Supplementary **Table 1:** Depicts KEGG annotation summary table of isolated *L. pentosus* strain (krglsrbmofpi2) from India.

**Supplementary Table 2:** Depicts geographical locations, sources, and genome sizes of *L. pentosus* strain worldwide

**Supplementary Table 3:** Classification of *L. pentosus* strain sources into animals, plants, humans, and others.

**Supplementary Table 4:** BPGA analysis results of 96 genomes of *L. pentosus* strains worldwide, which indicates a number of core genes, accessory genes, unique genes, and exclusively absent genes. The highlighted strain is the isolated strain used for the comparative study.

**Supplementary Table 5:** Represents bacteriocins present in 96 genomes of *L. pentosus* isolated from various geographical areas.

**Supplementary Table 6:** Represents the C-score, TM-score, and corresponding PDB hits for the predicted structures of three bacteriocins – Pentoplantaracin-EF, Pentopediocin, and Pentobovicin-as generated by I-TASSER.

**Supplementary Table 7:** Summary of binding interactions between bacteriocins and viral targets and their corresponding binding energies for each target-ligand pair.

---

**Fig. S1.**
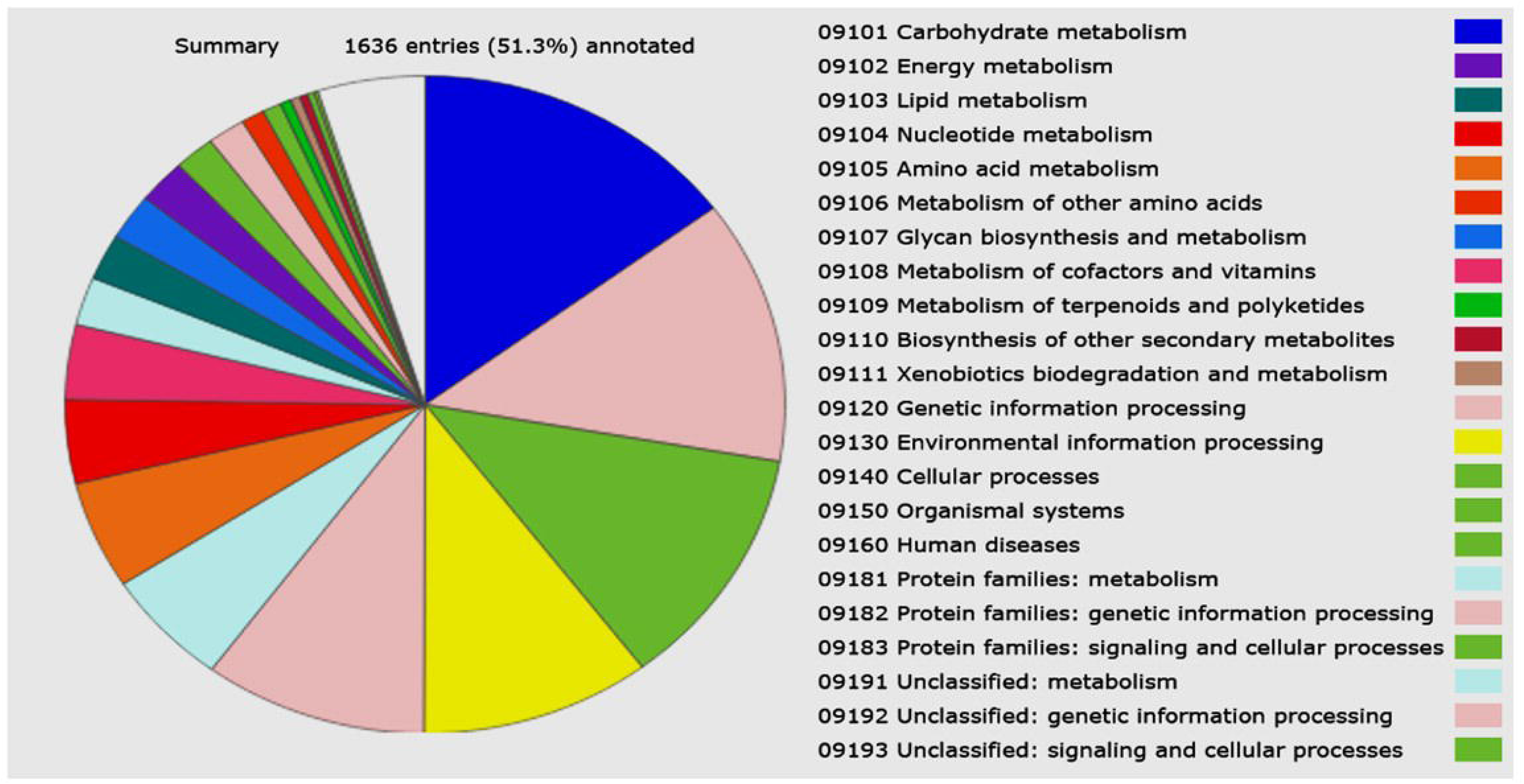
Depicts the summary of KEGG (BLAST KOALA) results of *L. pentosus* isolated from fermented foods of India

**Fig. S2.**
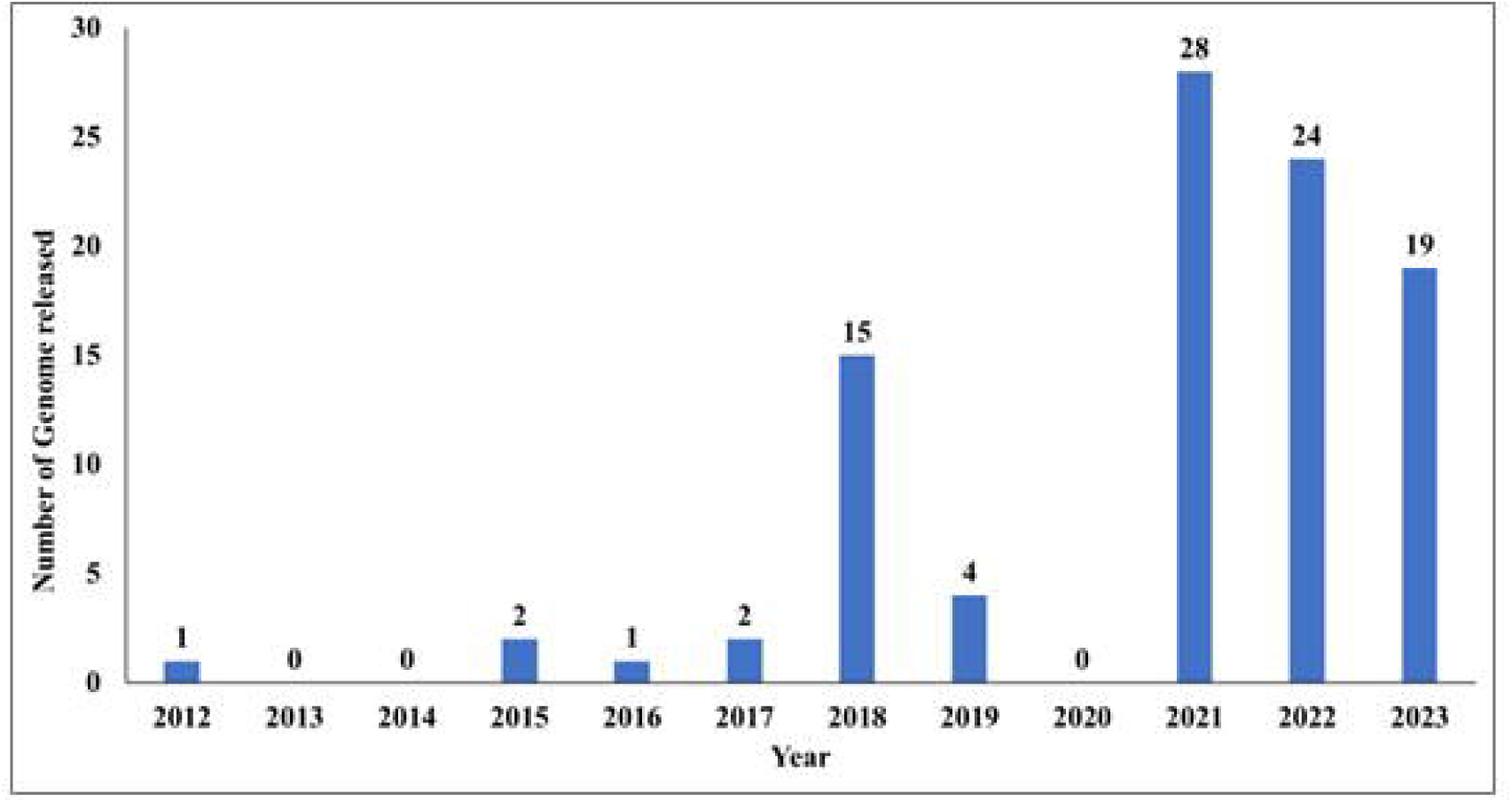

**Fig S3.**
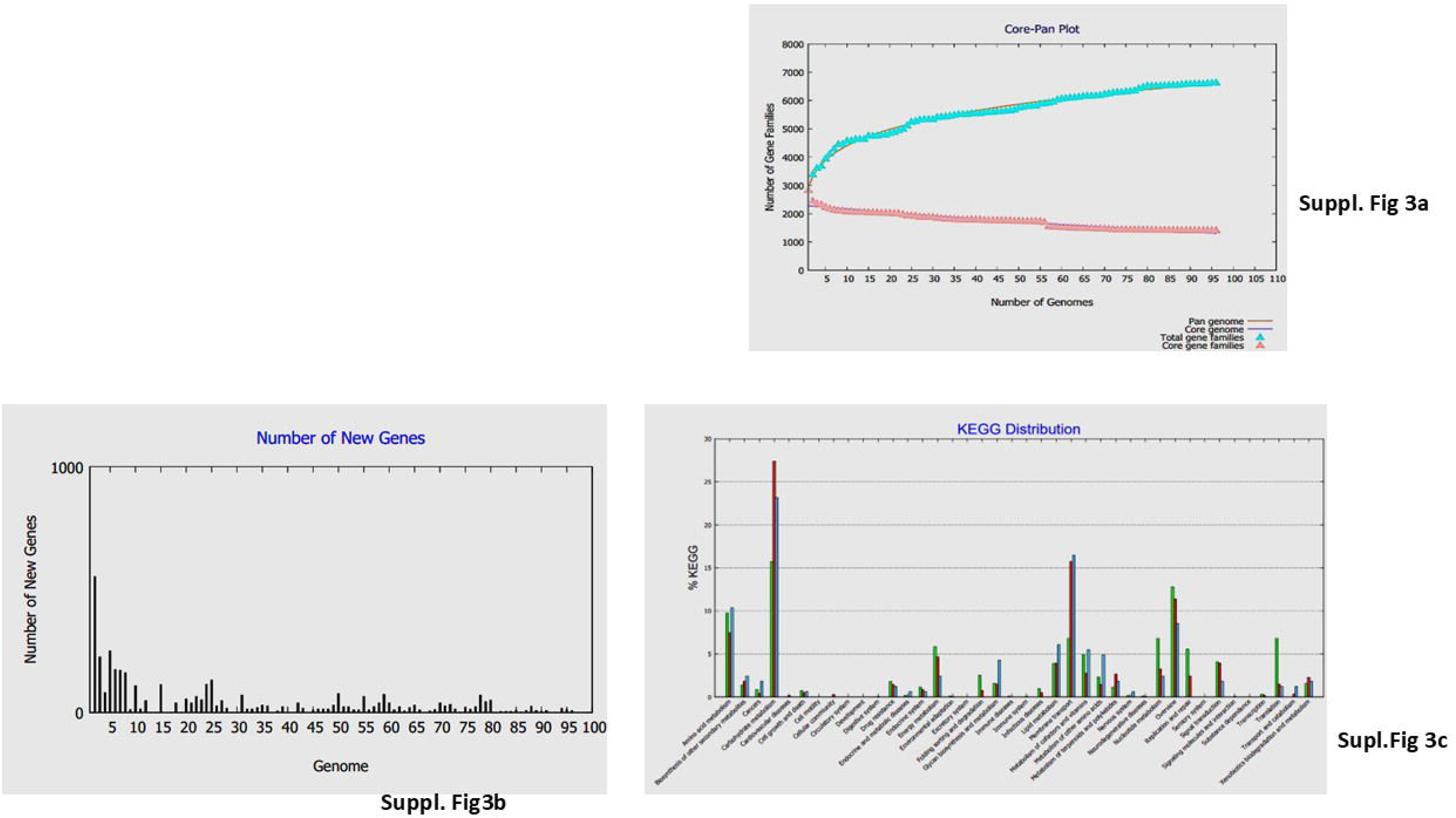
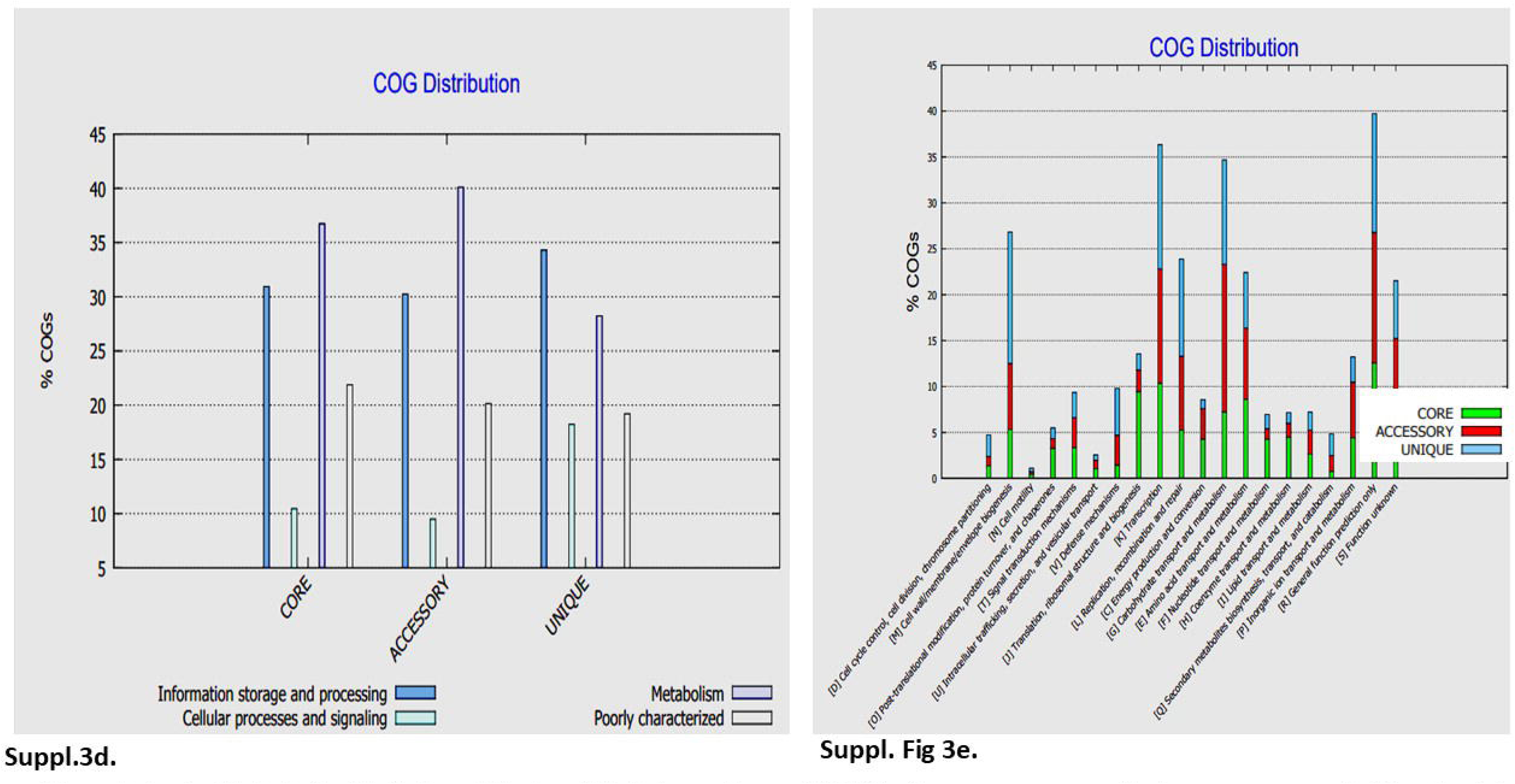
**a, b, c, d, e** Overview of BPGA analysis results, **a.** The core-pan plot of 96 strains of *L. pentosus* genomes, **b**. Illustrates the functional distributions of the core, accessory, and unique genes of *L. pentosus* genomes based on the KEGG pathway, **c. d** and **e** illustrate the distribution of Clusters of Orthologous Groups (COG) for the core, accessory, and unique genes among the 96 analyzed *L. pentosus* genomes (Results generated by the BPGA tool).

**Fig. S4.**
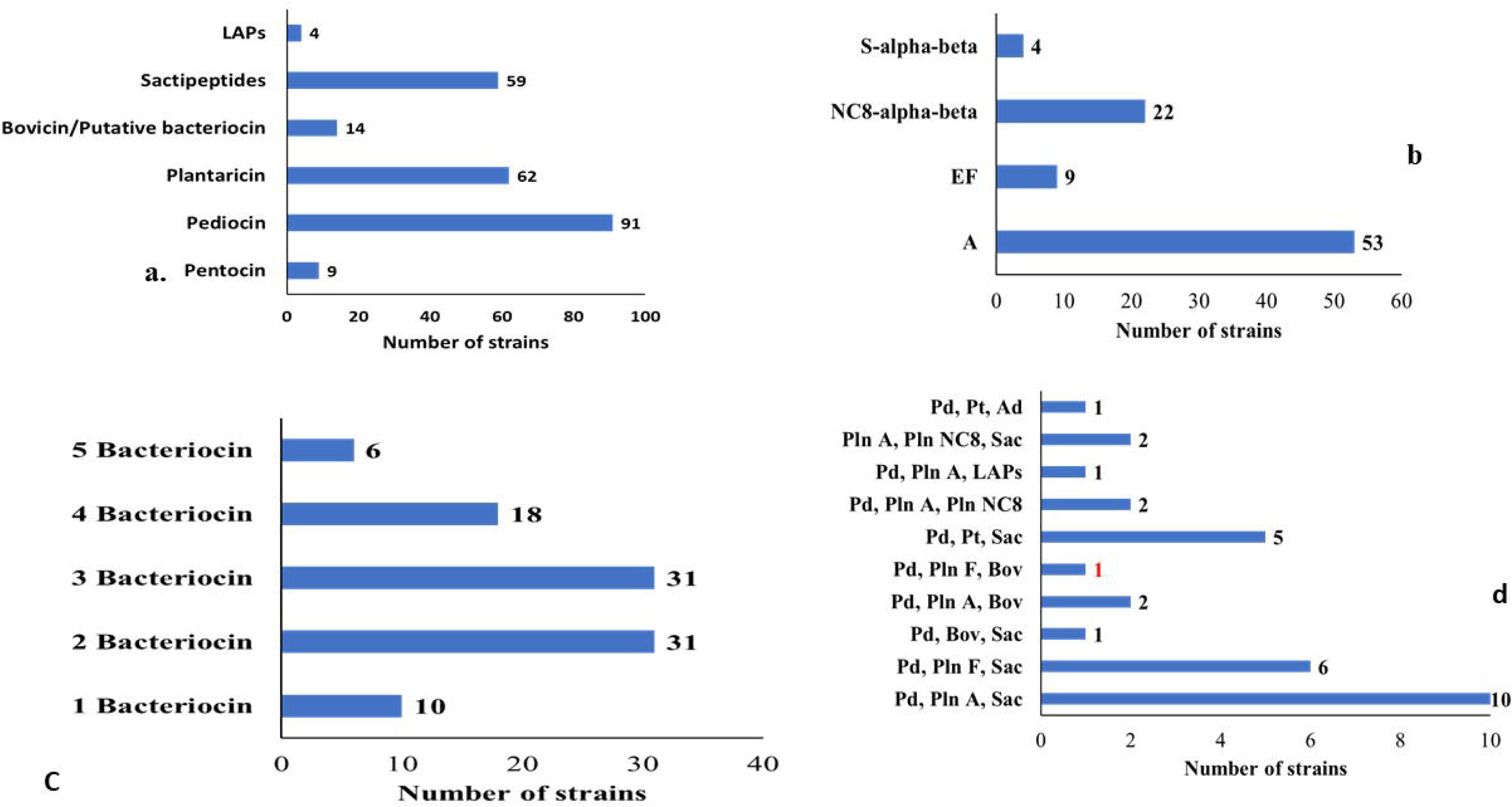
**a, b, c** Summarise the details of bacteriocins present in 96 genomes of *L. pentosus* and highlight the uniqueness of the Indian *L. pentosus* strain. **a.** Distribution of different types of bacteriocins across 96 *L. pentosus* genomes, illustrating the diversity present within this species; **b.** It shows different types of plantaricin among 62 strains with plantaracin. **c:** Summarise the details of bacteriocins present in 96 genomes of *L. pentosus* and highlight the uniqueness of the Indian *L. pentosus* strain. Overview of the number of strains producing varying quantities of bacteriocins, indicating the range of bacteriocin production within the analyzed strains; **d.** Unique combinations of three bacteriocins among 31 selected strains, with a focus on the distinct profile of the Indian strain (krglsrbmofpi2). The combination noted in red represents the Indian strain (krglsrbmofpi2). Pd—Pediocin; Pt—Pentocin; Ad—Acidocin; Pln—Plantaricin (including A and NC8 alpha-beta); Bov—Bovicin; LAPs—Linear Azole-Containing Peptides; Sac—Sactipeptides.

**Fig. S5.**
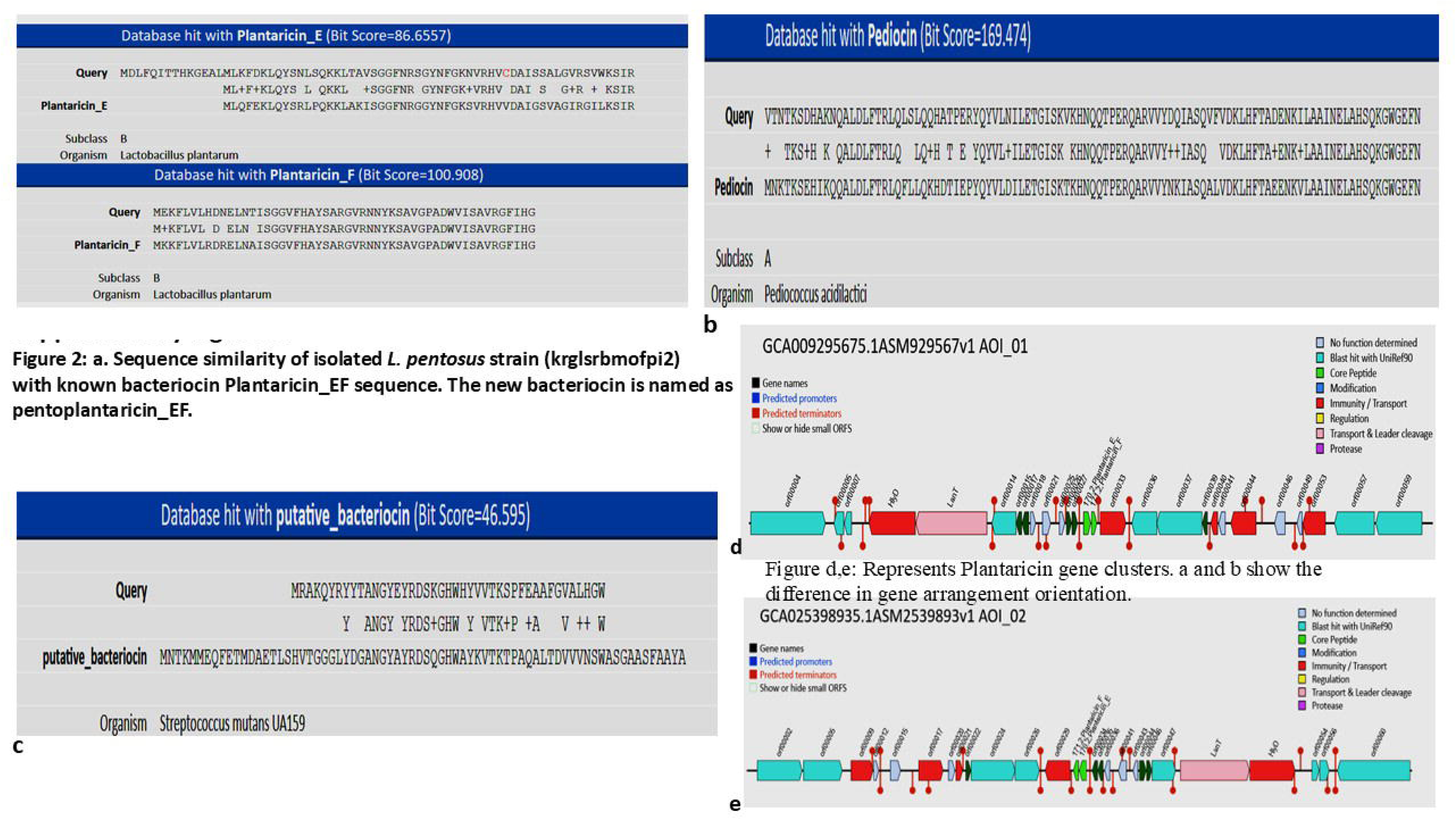

**Figure S6.**
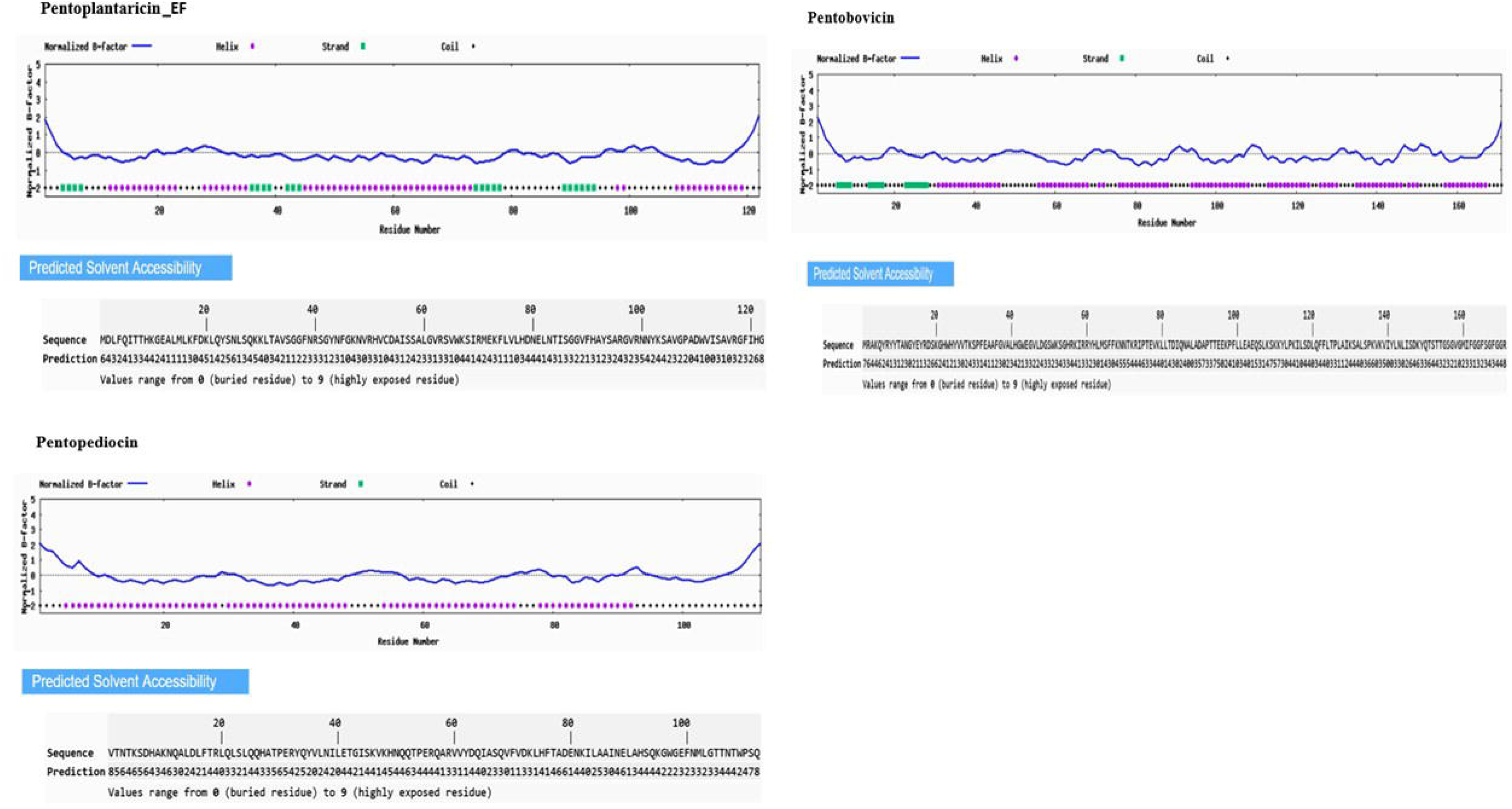
a, b, c: Predicted secondary structures and normalized B-factor indicating structural flexibility and solvent accessibility (I-TASSER) of Pentoplantaracin-EF, Pentobovicin, and Pentopediocin.

**Figure S7:**
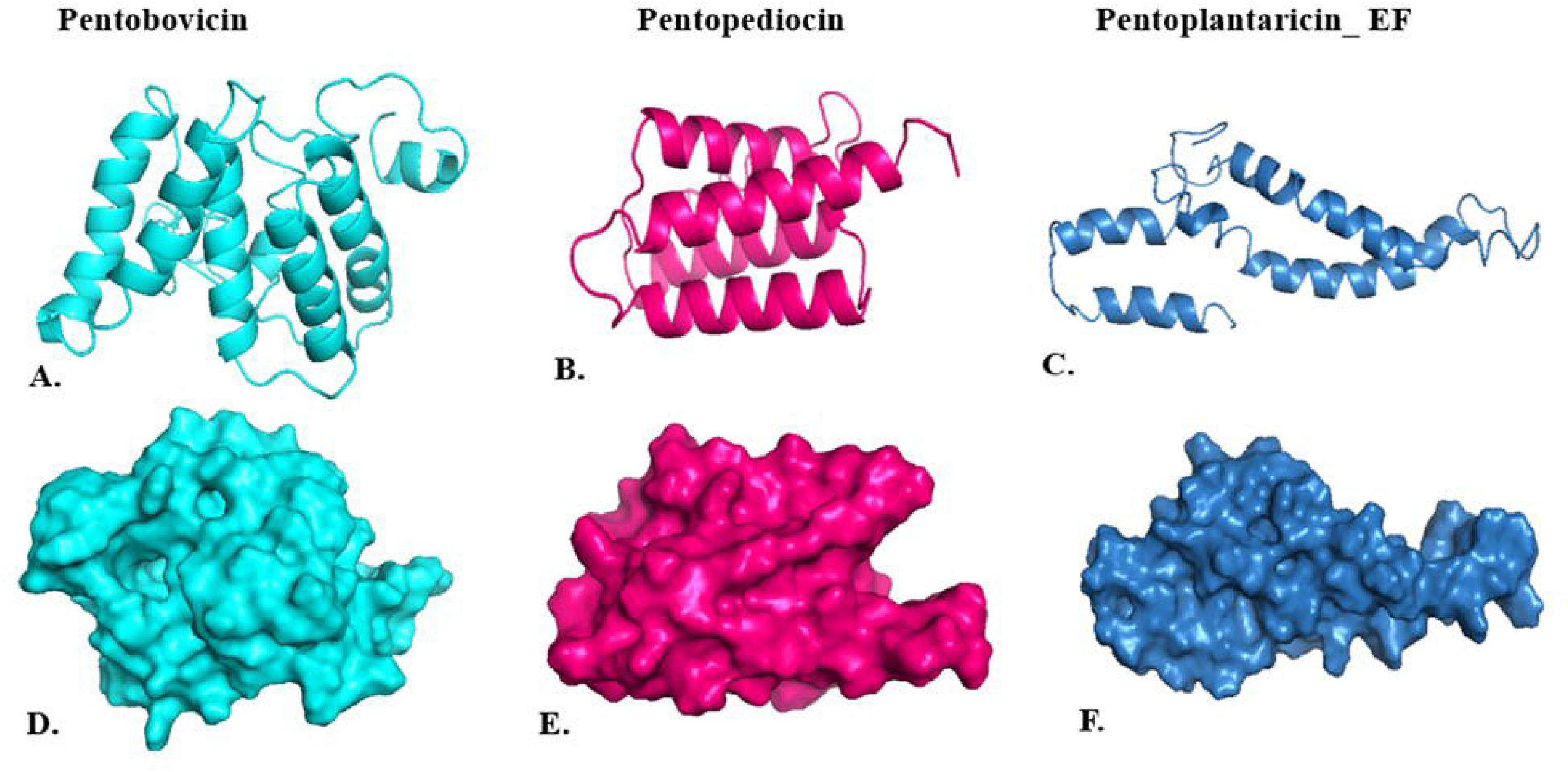
Represents the structure of the bacteriocins, Pentoplantaracin-EF, Pentobovicin, and Pentopediocin predicted by I-TASSER visualized with PyMOL. A, B, and C indicate cartoon structures, and D, E, and F for surface structures.

**Figure S8:**
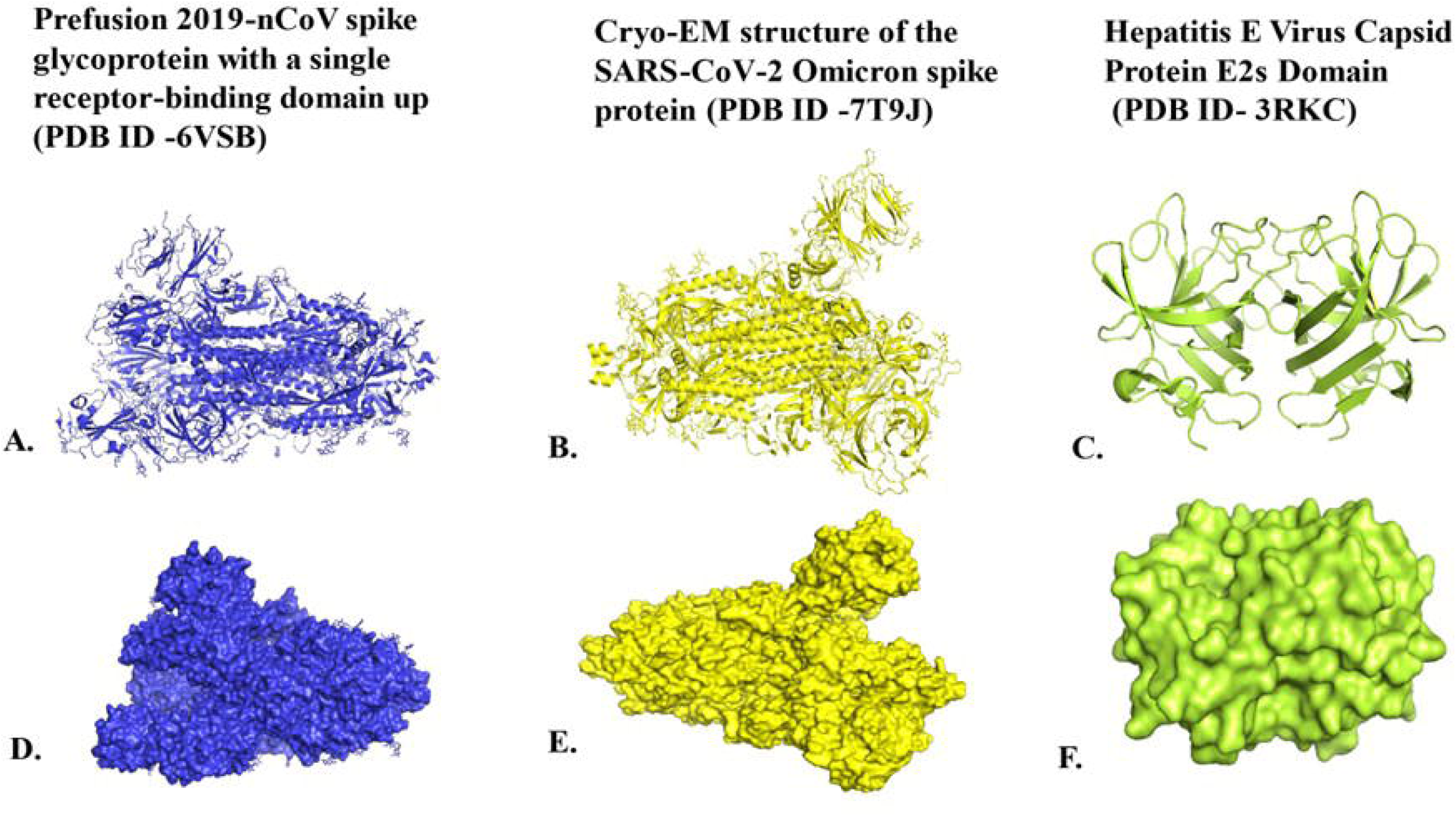
Target viral proteins visualized in PyMOL. A, B, and C indicate cartoon representations of the Prefusion 2019-nCov Spike glycoprotein (PDB ID:6VSB), the SARS-Cov-2 Omicron spike protein (PDB ID: 7T9J), and the Hepatitis E Virus Capsid Protein E2s domain (PDB ID: 3RKC). D, E, and F indicate corresponding surface representations.

## References

1. Abriouel H, Pérez Montoro B, Casado Muñoz MDC et al (2017) *In silico* genomic insights into aspects of food safety and defense mechanisms of a potentially probiotic *Lactobacillus pentosus* MP-10 isolated from brines of naturally fermented Aloreña green table olives. PLoS One 12(6): e0176801. 10.1371/journal.pone.0176801

2. Abriouel H, Pérez Montoro B, Casimiro-Soriguer CS et al (2017) Insight into potential probiotic markers predicted in *Lactobacillus pentosus* MP-10 genome sequence. Front Microbiol 8: 891. 10.3389/fmicb.2017.00891

3. Ahmed A, Siman-Tov G, Hall G et al (2019) Human antimicrobial peptides as therapeutics for viral infections. Viruses 11(8): 704. 10.3390/v11080704

4. Akman S, Akkoc S, Zeyrek CT et al (2023) Density functional modeling, and molecular docking with SARS-CoV-2 spike protein (Wuhan) and omicron S protein (variant) studies of new heterocyclic compounds including a pyrazoline nucleus. J Biomol Struct Dyn 41(22): 12951–12965. 10.1080/07391102.2023.2169765

5. Berman HM, Westbrook J, Feng Z et al (2000) The protein data bank. Nucleic Acids Res 28(1): 235–242. 10.1093/nar/28.1.235

6. Chaudhari NM, Gupta VK, Dutta C (2016) BPGA- an ultra-fast pan-genome analysis pipeline. Sci Rep 6: 24373. 10.1038/srep24373

7. Cheruvari A, Kammara R (2024) Draft genome sequence of a *Lactiplantibacillus pentosus* strain isolated from traditionally fermented rice. Access Microbiol 6(10): 000796.v3. 10.1099/acmi.0.000796.v3

8. Cheruvari A, Kammara R (2024) Genomic characterization and probiotic properties of *Lactiplantibacillus pentosus* isolated from fermented Rice. Probiotics Antimicrob Proteins 17(6): 4442–4464. 10.1007/s12602-024-10378-1

9. Cheruvari A, Kammara R (2024) Bacteriocins future perspectives: Substitutes to antibiotics. Food Control 168: 110834. 10.1016/j.foodcont.2024.110834

10. Collado MC, Sanz Y (2006) Method for direct selection of potentially probiotic Bifidobacterium strains from human feces based on their acid-adaptation ability. J Microbiol Methods 66(3): 560–563. 10.1016/j.mimet.2006.01.007

11. Collins FWJ, O’Connor PM, O’Sullivan O et al (2017) Bacteriocin gene-trait matching across the complete Lactobacillus pan-genome. Sci Rep 7(1): 3481. 10.1038/s41598-017-03339-y

12. Cotter PD, Hill C, Ross RP (2005) Bacteriocins: developing innate immunity for food. Nat Rev Microbiol 3(10): 777–788. 10.1038/nrmicro1273

13. Cotter PD, Ross RP, Hill C (2013) Bacteriocins - a viable alternative to antibiotics?. Nat Rev Microbiol 11(2): 95–105. 10.1038/nrmicro2937

14. Dassanayake MK, Khoo TJ, Chong CH, Di Martino P (2022) Molecular docking and *in-silico* analysis of natural biomolecules against dengue, ebola, zika, SARS-CoV-2 variants of concern and monkeypox virus. Int J Mol Sci 23(19): 11131. 10.3390/ijms231911131

15. de Jong A, van Hijum SA, Bijlsma JJ, et al (2006) BAGEL: a web-based bacteriocin genome mining tool. Nucleic Acids Res 34 (Web Server issue): W273–W279. 10.1093/nar/gkl237

16. Desta IT, Porter KA, Xia B et al (2020) Performance and its limits in rigid body protein-protein docking. Structure 28(9): 1071–1081.e3. 10.1016/j.str.2020.06.006

17. Ermolenko EI, Desheva YA, Kolobov AA et al (2019) Anti-influenza activity of Enterocin B in vitro and protective effect of bacteriocinogenic Enterococcal probiotic strain on influenza infection in mouse model. Probiotics Antimicrob Proteins 11(2): 705–712. 10.1007/s12602-018-9457-0

18. Evanovich E, Jeanne P, Mendonça DS, Guerreiro JF (2019) Comparative genomic analysis of *Lactobacillus plantarum*: An Overview. Int J Genomics 2019: 4973214. 10.1155/2019/4973214

19. Evivie SE, Huo GC, Igene JO, Bian X (2017) Some current applications, limitations and future perspectives of lactic acid bacteria as probiotics. Food Nutr Res 61(1): 1318034. 10.1080/16546628.2017.1318034

20. Fernandes J, Kumbhar R, Kulkarni R (2021) Bacteriocins from lactic acid bacteria: a natural strategy for inhibiting unwanted bacteria. Resonance 26(3): 387–398. 10.1007/s12045-021-1137-9

21. Franz CM, Huch M, Mathara JM et al (2014) African fermented foods and probiotics. Int J Food Microbiol 190:84–96. 10.1016/j.ijfoodmicro.2014.08.033

22. Gangaiah D, Mane SP, Tawari NR et al (2022) *In silico*, *in vitro* and *in vivo* safety evaluation of *Limosilactobacillus reuteri* strains ATCC PTA-126787 & ATCC PTA-126788 for potential probiotic applications. PLoS One 17(1): e0262663. 10.1371/journal.pone.0262663

23. Garcia-Gonzalez N, Bottacini F, van Sinderen D, et al (2022) Comparative genomics of *Lactiplantibacillus plantarum*: insights in to probiotic markers in strains isolated from the human gastrointestinal tract and fermented foods. Front Microbiol 13: 854266. 10.3389/fmicb.2022.854266

24. Goh YJ, Klaenhammer TR (2009) Genomic features of Lactobacillus species. Front Biosci (Landmark Ed) 14(4): 1362–1386. 10.2741/3313

25. Haque SkE, Bhadra S, Pal NK (2024) Exploring potential therapeutic candidates against COVID-19: a molecular docking study. Discov Mol 1: 5. 10.1007/s44345-024-00005-5

26. Jones G, Jindal A, Ghani U et al (2022) Elucidation of protein function using computational docking and hotspot analysis by ClusPro and FTMap. Acta Crystallogr D Struct Biol 78(Pt 6): 690–697. 10.1107/S2059798322002741

27. Kandasamy S, Yoo J, Yun J et al (2022) Probiogenomic *in-silico* analysis and safety Assessment of *Lactiplantibacillus plantarum* DJF10 strain isolated from Korean raw milk. Int J Mol Sci 23(22): 14494. 10.3390/ijms232214494

28. Kanehisa M, Sato Y, Morishima K (2016) BlastKOALA and GhostKOALA: KEGG tools for functional characterization of genome and metagenome sequences. J Mol Biol 428(4): 726–731. 10.1016/j.jmb.2015.11.006

29. Khelissa S, Chihib NE, Gharsallaoui A (2021) Conditions of nisin production by *Lactococcus lactis* subsp. *lactis* and its main uses as a food preservative. Arch Microbiol 203(2): 465–480. 10.1007/s00203-020-02054-z

30. Kozakov D, Beglov D, Bohnuud T et al (2013) How good is automated protein docking?. Proteins 81(12): 2159–2166. 10.1002/prot.24403

31. Kumariya R, Garsa AK, Rajput YS et al (2019) Bacteriocins: Classification, synthesis, mechanism of action and resistance development in food spoilage causing bacteria. Microb Pathog 128: 171–177. 10.1016/j.micpath.2019.01.002

32. Li SW, Chen YS, Lee YS et al (2017) Comparative genomic analysis of bacteriocin-producing *Weissella cibaria* 110. Appl Microbiol Biotechnol 101(3): 1227–1237. 10.1007/s00253-016-8073-8

33. Lalani S, Gew LT, Poh CL (2021) Antiviral peptides against *Enterovirus* A71 causing hand, foot and mouth disease. Peptides 136: 170443. 10.1016/j.peptides.2020.170443

34. Li Z, Song Q, Wang M et al (2021) Comparative genomics analysis of *Pediococcus acidilactici* species. J Microbiol 59(6): 573–583. 10.1007/s12275-021-0618-6

35. Liang CY, Yang CH, Lai CH et al (2019) Comparative genomics of 86 whole-genome Sequences in the six species of the *Elizabethkingia* genus reveals intraspecific and interspecific divergence. Sci Rep 9(1): 19167. 10.1038/s41598-019-55795-3

36. Lill MA, Danielson ML (2011) Computer-aided drug design platform using PyMOL. J Comput Aided Mol Des 25(1): 13–19. 10.1007/s10822-010-9395-8

37. Marco ML, Sanders ME, Gänzle M et al (2021) The International Scientific Association for Probiotics and Prebiotics (ISAPP) consensus statement on fermented foods. Nat Rev Gastroenterol Hepatol 18(3): 196–208. 10.1038/s41575-020-00390-5

38. Mashima I, Liao YC, Lin CH et al (2021) Comparative pan-genome analysis of oral Veillonella species. Microorganisms 9(8):1775. 10.3390/microorganisms9081775

39. Mukherjee A, Gómez-Sala B, O’Connor EM et al (2022) Global regulatory frameworks for fermented foods: a review. Front Nutr 9: 902642. 10.3389/fnut.2022.902642

40. Nuraida L (2015) A review: Health promoting lactic acid bacteria in traditional Indonesian fermented foods. Food Sci Hum Wellness 4(2): 47–55. 10.1016/j.fshw.2015.06.001

41. Page CA, Pérez-Díaz IM, Pan M, Barrangou R (2023) Genome-wide comparative analysis of *Lactiplantibacillus pentosus* isolates autochthonous to cucumber fermentation reveals subclades of divergent ancestry. Foods 12(13): 2455. 10.3390/foods12132455

42. Perales-Adán J, Rubiño S, Martínez-Bueno M et al (2018) LAB bacteriocins controlling the food isolated (drug-resistant) *Staphylococci*. Front Microbiol 9: 1143. 10.3389/fmicb.2018.01143

43. Quintero-Gil C, Parra-Suescún J, Lopez-Herrera A, Orduz S (2017) *In-silico* design and molecular docking evaluation of peptides derivatives from bacteriocins and porcine beta defensin-2 as inhibitors of Hepatitis E virus capsid protein. Virus disease 28(3): 281–288. 10.1007/s13337-017-0383-7

44. Quy PT, Bui T, Nguyen TM et al (2023) Combinatory *in silico* investigation for potential inhibitors from *Curcuma sahuynhensis* Škorničk. & N.S. Lý volatile phytoconstituents against influenza A hemagglutinin, SARS-CoV-2 main protease, and Omicron-variant spike protein. Open Chem 21(1): 20230109. 10.1515/chem-2023-0109

45. Rabah H, Silva S (2017) Applications of probiotic bacteria and dairy foods in health. In: Curr Res Microbiol. Open Access eBooks, Wilmington, USA

46. Rajagopal K (2013) Draft genome sequence of *Enterococcus raffinosus* strain CFTRI 2200, isolated from infant fecal material. Genome Announc 1(6): e00932–13. 10.1128/genomeA.00932-13

47. Satish Kumar R, Kanmani P, Yuvaraj N et al (2013) Traditional Indian fermented foods: a rich source of lactic acid bacteria. Int J Food Sci Nutr 64(4): 415–428. 10.3109/09637486.2012.746288

48. Seeliger D, de Groot BL (2010) Ligand docking and binding site analysis with PyMOL and Autodock/Vina. J Comput Aided Mol Des 24(5): 417–422. 10.1007/s10822-010-9352-6

49. Stergiou OS, Tegopoulos K, Kiousi DE, et al (2021) Whole-genome sequencing, phylogenetic and genomic analysis of *Lactiplantibacillus pentosus* L33, a potential probiotic strain isolated from fermented sausages. Front Microbiol 12: 746659. 10.3389/fmicb.2021.746659

50. Surve S, Shinde DB, Kulkarni R (2022) Isolation, characterization and comparative genomics of potentially probiotic *Lactiplantibacillus plantarum* strains from Indian foods. Sci Rep 12(1): 1940. 10.1038/s41598-022-05850-3

51. Teuber M (2008) Lactic acid bacteria. In: Biotech: second, completely revised edition, vol 1–12. Wiley-VCH, Weinheim. 10.1002/9783527620999.ch10

52. Vajda S, Yueh C, Beglov D et al (2017) New additions to the ClusPro server motivated by CAPRI. Proteins 85(3): 435–444. 10.1002/prot.25219

53. van Heel AJ, de Jong A, Song C, et al (2018) BAGEL4: a user-friendly web server to thoroughly mine RiPPs and bacteriocins. Nucleic Acids Res 46(W1): W278–W281. 10.1093/nar/gky383

54. Vijayalakshmi S, Adeyemi D, Choi IY et al (2020) Comprehensive *in silico* analysis of lactic acid bacteria for the selection of desirable probiotics. LWT 130(5): 109617. 10.1016/j.lwt.2020.109617

55. Wang S, Yang B, Ross RP et al (2020) Comparative genomics analysis of *Lactobacillus ruminis* from different niches. Genes (Basel) 11(1): 70. 10.3390/genes11010070

56. Webber JL, Namivandi-Zangeneh R, Drozdek S et al (2021) Incorporation and antimicrobial activity of nisin Z within carrageenan/chitosan multilayers. Sci Rep 11(1): 1690. 10.1038/s41598-020-79702-3

57. Yang J, Zhang Y (2015) I-TASSER server: new development for protein structure and function predictions. Nucleic Acids Res 43(W1): W174–W181. 10.1093/nar/gkv342

58. Ye K, Li P, Gu Q (2020) Complete genome sequence analysis of a strain *Lactobacillus pentosus* ZFM94 and its probiotic characteristics. Genomics 112(5): 3142–3149. 10.1016/j.ygeno.2020.05.015

59. Goh YJ, Klaenhammer TR (2009) Genomic features of Lactobacillus species. Front Biosci (Landmark Ed) 14(4): 1362–1386. 10.2741/3313

60. Yuan S, Chan HCS, Hu Z (2017) Using PyMOL as a platform for computational drug design. Wiley Interdiscip Rev Comput Mol Sci 7(2): e1298. 10.1002/wcms.1298

61. Zarzosa-Moreno D, Avalos-Gómez C, Ramírez-Texcalco LS et al (2020) Lactoferrin and Its derived peptides: an alternative for combating virulence mechanisms developed by pathogens. Molecules 25(24): 5763. 10.3390/molecules25245763

62. Zheng W, Zhang C, Li Y et al (2021) Folding non-homologous proteins by coupling deep-learning contact maps with I-TASSER assembly simulations. Cell Rep Methods 1(3): 100014. 10.1016/j.crmeth.2021.100014

63. Zhou X, Zheng W, Li Y et al (2022) I-TASSER-MTD: a deep-learning-based platform for multi-domain protein structure and function prediction. Nat Protoc 17(10): 2326–2353. 10.1038/s41596-022-00728-0

64. Zimina M, Babich O, Prosekov A et al (2020) Overview of global trends in classification, methods of preparation and application of bacteriocins. Antibiotics (Basel) 9(9): 553. 10.3390/antibiotics9090553

