## Supplementary table 1 for "Bacteriocin Diversity and Antiviral Potential of *Lactiplantibacillus pentosus from* Fermented Rice"

**Supplementary Table 1: Depicts KEGG annotation summary table of isolated L. *pentosus* strain (krglsrbmofpi2) from India.**

| Sl No. | Global and overview maps | No. of Entries |
| --- | --- | --- |
| 1 | Carbohydrate metabolism | 249 |
| 2 | Protein families: genetic information processing | 214 |
| 3 | Protein families: signaling and cellular processes | 196 |
| 4 | Environmental information | 170 |
| 5 | Genetic information processing | 165 |
| 6 | Unclassified: metabolism | 95 |
| 7 | Amino acid metabolism | 87 |
| 8 | Nucleotide metabolism | 68 |
| 9 | Metabolism of cofactors and vitamins | 61 |
| 10 | Protein families: metabolism | 40 |
| 11 | Glycan biosynthesis and metabolism | 37 |
| 12 | Lipid metabolism | 36 |
| 13 | Energy metabolism | 36 |
| 14 | Unclassified: signaling and cellular processes | 31 |
| 15 | Unclassified: genetic information processing | 28 |
| 16 | Metabolism of other amino acids | 17 |
| 17 | Cellular processes | 15 |
| 18 | Metabolism of terpenoids and polyketides | 8 |
| 19 | Xenobiotics biodegradation and metabolism | 7 |
| 20 | Biosynthesis of other secondary metabolites | 6 |
| 21 | Organismal systems | 5 |
| 22 | Human diseases | 3 |
| 23 | Unclassified | 79 |
