## Supplementary table 2 for "Bacteriocin Diversity and Antiviral Potential of *Lactiplantibacillus pentosus from* Fermented Rice"

| **Sl no.** | **Genome accession No.** | **Genome location/ submitter location** | **Source of Sample** | **Genome size** | **Genome Release date** |
| --- | --- | --- | --- | --- | --- |
| 1. | GCA_000271445.1 | Nigeria | vagina of a healthy Nigerian woman | 3.4 Mb | June 2012 |
| 2. | GCA_001188985.1 | China | Temperate deciduous forest biome soil | 3.7 Mb | July 2015 |
| 3. | GCA_001433755.1 | Shanghai Majorbio, China | Corn silage | 3.6 Mb | November 2015 |
| 4. | GCA_032195525.1 | USA: Chaska, MN | cucumber fermentation brine | 3.9 Mb | July 2016 |
| 5. | GCA_002211885.1 | Taiwan: Meinong District, Kaohsiung | mustard pickles | 3.6 Mb | July 2017 |
| 6. | GCA_002751855.1 | Switzerland. | milk product | 3.7 Mb | November 2017 |
| 7. | GCA_002850015.1 | USA: Davis, CA | olive fermentation | 3.7 Mb | January 2018 |
| 8. | GCA_002993385.1 | Spain | brines of olives | 3.8 Mb | March 2018 |
| 9. | GCA_002993395.1 | Spain | brines of olives | 3.8 Mb | March 2018 |
| 10. | GCA_002993425.1 | Spain | brines of olives | 3.9 Mb | March 2018 |
| 11. | GCA_002993435.1 | Spain | brines of olives | 3.8 Mb | March 2018 |
| 12. | GCA_002993465.1 | Spain | brines of olives | 4 Mb | March 2018 |
| 13. | GCA_002993485.1 | Spain | brines of olives | 3.9 Mb | March 2018 |
| 14. | GCA_003627295.1 | China: Tangbiao village Lanxi town Zhejiang province | Fermented vegetables | 3.7 Mb | October 2018 |
| 15. | GCA_003627375.1 | China: Zhejiang Maternal and Child Health Hospital | healthy infant fecal samples | 3.7 Mb | October 2018 |
| 16. | GCA_003641185.1 | Korea University | DSM 20314, type strain | 3.7 Mb | October 2018 |
| 17. | GCA_003702565.1 | Spain | biofilm | 3.8 Mb | October 2018 |
| 18. | GCA_003702605.1 | Spain | biofilm | 3.8 Mb | October 2018 |
| 19. | GCA_003702625.1 | Spain | biofilm | 3.8 Mb | October 2018 |
| 20. | GCA_003702635.1 | Spain | biofilm | 3.8 Mb | October 2018 |
| 21. | GCA_003702665.1 | Spain | biofilm | 3.8 Mb | October 2018 |
| 22. | GCA_004354685.1 | Denmark | ATCC:8041 | 3.7 Mb | March 2019 |
| 23. | GCA_007991835.1 | NBRC, Tokyo, Japan | Corn silage | 3.5 Mb | July 2019 |
| 24. | GCA_009295675.1 | India | Fermented rice | 3.7 Mb | October 2019 |
| 25. | GCA_009812415.1 | China: Dazhou | pickle | 4 Mb | December 2019 |
| 26. | GCA_016804305.1 | China: Meishan | Pickle | 3.8 Mb | February 2021 |
| 27. | GCA_017742275.1 | Taiwan | Pickled cucumber | 3.7 Mb | April 2021 |
| 28. | GCA_018069575.1 | China: Sichuan Province | paocai water | 3.8 Mb | April 2021 |
| 29. | GCA_018403455.1 | Gifu University, Japan | Awa-bancha | 3.7 Mb | April 2021 |
| 30. | GCA_018982875.1 | Italy: Potenza | olive brine | 3.9 Mb | June 2021 |
| 31. | GCA_018991285.1 | USA: Mount Olive, NC | cucumber fermentation brine | 3.6 Mb | June 2021 |
| 32. | GCA_018991345.1 | USA: Mount Olive, NC | cucumber fermentation brine | 3.7 Mb | June 2021 |
| 33. | GCA_018991355.1 | USA: Mount Olive, NC | cucumber fermentation brine | 3.6 Mb | June 2021 |
| 34. | GCA_018991365.1 | USA: Mount Olive, NC | cucumber fermentation brine | 3.7 Mb | June 2021 |
| 35. | GCA_018991375.1 | USA: Mount Olive, NC | cucumber fermentation brine | 3.7 Mb | June 2021 |
| 36. | GCA_018991425.1 | USA: Mount Olive, NC | cucumber fermentation brine | 3.7 Mb | June 2021 |
| 37. | GCA_018991465.1 | USA: Mount Olive, NC | cucumber fermentation brine | 3.8 Mb | June 2021 |
| 38. | GCA_018991535.1 | USA: Chaska, MN | cucumber fermentation brine | 3.8 Mb | June 2021 |
| 39. | GCA_018991565.1 | USA: Mount Olive, NC | cucumber fermentation brine | 3.7 Mb | June 2021 |
| 40. | GCA_018993275.1 | USA: Mount Olive, NC | cucumber fermentation brine | 3.8 Mb | June 2021 |
| 41. | GCA_018993345.1 | USA: Mount Olive, NC | cucumber fermentation brine | 3.7 Mb | June 2021 |
| 42. | GCA_018993445.1 | USA: Mount Olive, NC | cucumber fermentation brine | 3.7 Mb | June 2021 |
| 43. | GCA_018993485.1 | USA: Mount Olive, NC | cucumber fermentation brine | 3.7 Mb | June 2021 |
| 44. | GCA_018993495.1 | USA: Mount Olive, NC | cucumber fermentation brine | 3.7 Mb | June 2021 |
| 45. | GCA_018993525.1 | USA: Mount Olive, NC | cucumber fermentation brine | 3.8 Mb | June 2021 |
| 46. | GCA_018993585.1 | USA: Mount Olive, NC | cucumber fermentation brine | 3.8 Mb | June 2021 |
| 47. | GCA_018993625.1 | USA: Mount Olive, NC | cucumber fermentation brine | 3.8 Mb | June 2021 |
| 48. | GCA_018993635.1 | USA: Mount Olive, NC | cucumber fermentation brine | 3.7 Mb | June 2021 |
| 49. | GCA_018993665.1 | USA: Chaska, MN | cucumber fermentation brine | 3.8 Mb | June 2021 |
| 50. | GCA_018993725.1 | USA: Mount Olive, NC | cucumber fermentation brine | 3.7 Mb | June 2021 |
| 51. | GCA_018993945.1 | USA: Mount Olive, NC | cucumber fermentation brine | 3.6 Mb | June 2021 |
| 52. | GCA_020181715.1 | Greece: Athens | meat samples | 3.9 Mb | September 2021 |
| 53. | GCA_020532005.1 | Thailand: Bangkok | fermented fish (Pla-som) | 3.9 Mb | October 2021 |
| 54. | GCA_021384425.1 | USDA-ARS, USA | Cucumber fermentation tank | 3.7 Mb | Jan 2022 |
| 55. | GCA_022701335.1 | China: Chongqing | Pickle | 3.9 Mb | March 2022 |
| 56. | GCA_022936785.1 | Italy: Potenza | fermented table olives | 3.9 Mb | April 2022 |
| 57. | GCA_022936805.1 | Italy: Potenza | olives brine | 3.7 Mb | April 2022 |
| 58. | GCA_022936825.1 | Italy: Potenza | fermented table olives | 3.9 Mb | April 2022 |
| 59. | GCA_023823145.1 | Thailand | pickled weed | 3.8 Mb | June 2022 |
| 60. | GCA_023972895.1 | Norway | Olives | 3.9 Mb | June 2022 |
| 61. | GCA_023980805.1 | Norway | Olives | 3.8 Mb | June 2022 |
| 62. | GCA_025122375.1 | France | wheat sourdough | 3.7 Mb | September 2022 |
| 63. | GCA_025129205.1 | Switzerland | milk products | 3.7 Mb | September 2022 |
| 64. | GCA_025188445.1 | Egypt | cheese (Domiati) | 3.8 Mb | September 2022 |
| 65. | GCA_025190245.1 | Poland | artisanal fermented pickle | 3.7 Mb | September 2022 |
| 66. | GCA_025190265.1 | Thailand | fermented tea leaf | 3.6 Mb | September 2022 |
| 67. | GCA_025190285.1 | Thailand | fish cake | 3.6 Mb | September 2022 |
| 68. | GCA_025190295.1 | Thailand | pickle (saumure) | 3.8 Mb | September 2022 |
| 69. | GCA_025190325.1 | Thailand | fermented bamboo | 3.7 Mb | September 2022 |
| 70. | GCA_025190345.1 | Thailand | fermented fish | 3.8 Mb | September 2022 |
| 71. | GCA_025190365.1 | unknown, Brigham Young University, United States | corn silage | 3.6 Mb | September 2022 |
| 72. | GCA_025190375.1 | Thailand | pork sausage | 3.5 Mb | September 2022 |
| 73. | GCA_025190395.1 | Spain | olive fermentation | 3.9 Mb | September 2022 |
| 74. | GCA_025190425.1 | Spain | olive fermentation | 3.7 Mb | September 2022 |
| 75. | GCA_025190445.1 | Egypt | cheese (Domiati) | 3.8 Mb | September 2022 |
| 76. | GCA_025190465.1 | Spain | olive fermentation | 3.8 Mb | September 2022 |
| 77. | GCA_025398935.1 | China: Xi'an | Jiangshui | 3.6 Mb | November 2022 |
| 78. | GCA_026222675.1 | South Korea | Human Milk | 3.6 Mb | January 2023 |
| 79. | GCA_027675785.1 | China: Shenzhen | fecal material, Homo sapiens | 3.7 Mb | January 2023 |
| 80. | GCA_027675885.1 | China: Shenzhen | fecal material, Homo sapiens | 3.7 Mb | January 2023 |
| 81. | GCA_027676485.1 | China: Shenzhen | fecal material, Homo sapiens | 3.7 Mb | January 2023 |
| 82. | GCA_028464285.1 | China: Chongqing | fermented bamboo shoots | 3.6 Mb | February 2023 |
| 83. | GCA_029228745.1 | China: Beijing | Chinese traditional pickle | 3.9 Mb | March 2023 |
| 84. | GCA_029542285.1 | China | phyllo sphere epiphyte | 3.6 Mb | April 2023 |
| 85. | GCA_029813085.1 | China: Yunnan | vegetable | 3.7 Mb | April 2023 |
| 86. | GCA_030480485.1 | China: Nanjing | Dairy | 3.8 Mb | July 2023 |
| 87. | GCA_030489685.1 | China: Nanjing | Dairy | 3.7 Mb | July 2023 |
| 88. | GCF_030549395.1 | China | fermented food | 3.8 Mb | July 2023 |
| 89. | GCA_900092635.1 | University of Jaén, Spain | Fermented Alorena table olives | 3.9 Mb | July 2023 |
| 90. | GCA_030578355.1 | Thailand: Nakhon Pathom | pickled onion | 3.7 Mb | August 2023 |
| 91. | GCA_032190575.1 | USA: Mount Olive, NC | cucumber fermentation brine | 3.9 Mb | September 2023 |
| 92. | GCA_032190655.1 | USA: Mount Olive, NC | cucumber fermentation brine | 4.1 Mb | September 2023 |
| 93. | GCA_032190675.1 | USA: Mount Olive, NC | cucumber fermentation brine | 4 Mb | September 2023 |
| 94. | GCA_032190695.1 | USA: Mount Olive, NC | cucumber fermentation brine | 3.7 Mb | September 2023 |
| 95. | GCA_032190735.1 | USA: Mount Olive, NC | cucumber fermentation brine | 3.9 Mb | September 2023 |
| 96. | GCA_018993965.2 | USA: Mount Olive, NC | cucumber fermentation brine | 3.9 Mb | December 2023 |

**Suppl.Table 2: Depicts geographical locations, sources, and genome sizes of *L. pentosus* strain worldwide.**
