## Supplementary table 3 for "Bacteriocin Diversity and Antiviral Potential of *Lactiplantibacillus pentosus from* Fermented Rice"

| **Sl No.** | **Sources of *L. pentosus*** | **No. of Samples** |
| --- | --- | --- |
| 1. | **Human**  Vagina of a healthy Nigerian woman, Human Milk, Fecal material. | 7 |
| 2. | **Plant**  Corn silage, Vegetable Pickles, Mustard Pickles, olives, Fermented rice, Awa-bancha (Japanese dark tea), pickled weed, wheat sourdough, fermented tea leaf, fermented bamboo, phyllosphere epiphyte(alfalfa). | 65 |
| 3. | **Animal**  Milk products, cheese (Domiati), pork sausage, fermented fish, meat samples, fish cake. | 10 |
| 4. | **Other samples**   a. **Environmental –** Soil. b. **Other food items-** Jiangshui, Chinese traditional pickle, artisanal fermented pickle, paocai water, pickle (saumure - salt water). c. **Biofilm** d. **Unknown source-** Microbial/viral/environmental/type culture | 1  6  5  2 |
|  | **Total samples** | **96** |

**Suppl Table 3: Classification of *L. pentosus* strain sources into animals, plants, humans, and others.**
