## Supplementary table 4 for "Bacteriocin Diversity and Antiviral Potential of *Lactiplantibacillus pentosus from* Fermented Rice"

| **Genome No.** | **NCBI Accession**  **No.** | **No. of core genes** | **No. of accessory genes** | **No. of unique genes** | **No. of exclusively absent genes** |
| --- | --- | --- | --- | --- | --- |
| 1 | GCA_000271445.1.faa | 1420 | 1360 | 62 | 20 |
| 2 | GCA_001433755.1.faa | 1420 | 1569 | 17 | 12 |
| 3 | GCA_002211885.1.faa | 1420 | 1487 | 20 | 18 |
| 4 | GCA_002751855.1.faa | 1420 | 1628 | 0 | 3 |
| 5 | GCA_002850015.1.faa | 1420 | 1489 | 7 | 12 |
| 6 | GCA_002993385.1.faa | 1420 | 1642 | 33 | 11 |
| 7 | GCA_002993395.1.faa | 1420 | 1619 | 23 | 12 |
| 8 | GCA_002993425.1.faa | 1420 | 1779 | 15 | 0 |
| 9 | GCA_002993435.1.faa | 1420 | 1668 | 0 | 0 |
| 10 | GCA_002993465.1.faa | 1420 | 1786 | 20 | 9 |
| 11 | GCA_002993485.1.faa | 1420 | 1741 | 5 | 1 |
| 12 | GCA_003627295.1.faa | 1420 | 1546 | 0 | 0 |
| 13 | GCA_003627375.1.faa | 1420 | 1544 | 1 | 0 |
| 14 | GCA_003641185.1.faa | 1420 | 1597 | 1 | 0 |
| 15 | GCA_003702565.1.faa | 1420 | 1705 | 0 | 0 |
| 16 | GCA_003702605.1.faa | 1420 | 1709 | 0 | 0 |
| 17 | GCA_003702625.1.faa | 1420 | 1712 | 0 | 0 |
| 18 | GCA_003702635.1.faa | 1420 | 1663 | 20 | 10 |
| 19 | GCA_003702665.1.faa | 1420 | 1707 | 1 | 0 |
| 20 | GCA_004354685.1.faa | 1420 | 1665 | 29 | 0 |
| 21 | GCA_007991835.1.faa | 1420 | 1653 | 26 | 1 |
| 22 | GCA_009295675.1.faa | 1420 | 1588 | 25 | 0 |
| 23 | GCA_017742275.1.faa | 1420 | 1577 | 11 | 8 |
| 24 | GCA_018069575.1.faa | 1420 | 1658 | 12 | 9 |
| 25 | GCA_018403455.1.faa | 1420 | 1627 | 78 | 0 |
| 26 | GCA_018982875.1.faa | 1420 | 1760 | 1 | 0 |
| 27 | GCA_018991285.1.faa | 1420 | 1541 | 3 | 0 |
| 28 | GCA_018991345.1.faa | 1420 | 1573 | 0 | 0 |
| 29 | GCA_018991355.1.faa | 1420 | 1556 | 0 | 0 |
| 30 | GCA_018991365.1.faa | 1420 | 1570 | 0 | 0 |
| 31 | GCA_018991375.1.faa | 1420 | 1605 | 21 | 12 |
| 32 | GCA_018991425.1.faa | 1420 | 1636 | 7 | 4 |
| 33 | GCA_018991465.1.faa | 1420 | 1690 | 0 | 0 |
| 34 | GCA_018991535.1.faa | 1420 | 1662 | 2 | 2 |
| 35 | GCA_018991565.1.faa | 1420 | 1590 | 0 | 0 |
| 36 | GCA_018993275.1.faa | 1420 | 1708 | 0 | 0 |
| 37 | GCA_018993345.1.faa | 1420 | 1582 | 0 | 0 |
| 38 | GCA_018993445.1.faa | 1420 | 1629 | 0 | 0 |
| 39 | GCA_018993485.1.faa | 1420 | 1598 | 15 | 1 |
| 40 | GCA_018993495.1.faa | 1420 | 1552 | 0 | 0 |
| 41 | GCA_018993525.1.faa | 1420 | 1709 | 0 | 1 |
| 42 | GCA_018993585.1.faa | 1420 | 1645 | 10 | 7 |
| 43 | GCA_018993625.1.faa | 1420 | 1674 | 1 | 1 |
| 44 | GCA_018993635.1.faa | 1420 | 1594 | 0 | 0 |
| 45 | GCA_018993665.1.faa | 1420 | 1706 | 10 | 1 |
| 46 | GCA_018993725.1.faa | 1420 | 1607 | 13 | 2 |
| 47 | GCA_018993945.1.faa | 1420 | 1534 | 0 | 1 |
| 48 | GCA_018993965.2.faa | 1420 | 1697 | 4 | 3 |
| 49 | GCA_020181715.1.faa | 1420 | 1664 | 13 | 5 |
| 50 | GCA_020532005.1_.faa | 1420 | 1764 | 36 | 0 |
| 51 | GCA_021384425.1.faa | 1420 | 1617 | 10 | 3 |
| 52 | GCA_022936785.1.faa | 1420 | 1721 | 12 | 0 |
| 53 | GCA_022936805.1_.faa | 1420 | 1586 | 9 | 0 |
| 54 | GCA_022936825.1.faa | 1420 | 1802 | 4 | 1 |
| 55 | GCA_023823145.1.faa | 1420 | 1615 | 59 | 3 |
| 56 | GCA_023972895.1.faa | 1420 | 1655 | 5 | 15 |
| 57 | GCA_023980805.1.faa | 1420 | 1344 | 22 | 131 |
| 58 | GCA_025122375.1.faa | 1420 | 1590 | 28 | 6 |
| 59 | GCA_025129205.1.faa | 1420 | 1524 | 61 | 5 |
| 60 | GCA_025188445.1.faa | 1420 | 1669 | 35 | 9 |
| 61 | GCA_025190245.1.faa | 1420 | 1629 | 9 | 3 |
| 62 | GCA_025190265.1.faa | 1420 | 1567 | 18 | 3 |
| 63 | GCA_025190285.1.faa | 1420 | 1564 | 5 | 8 |
| 64 | GCA_025190295.1.faa | 1420 | 1614 | 18 | 8 |
| 65 | GCA_025190325.1.faa | 1420 | 1627 | 24 | 4 |
| 66 | GCA_025190345.1.faa | 1420 | 1682 | 7 | 5 |
| 67 | GCA_025190365.1.faa | 1420 | 1599 | 0 | 0 |
| 68 | GCA_025190375.1.faa | 1420 | 1512 | 3 | 5 |
| 69 | GCA_025190395.1.faa | 1420 | 1766 | 3 | 1 |
| 70 | GCA_025190425.1.faa | 1420 | 1594 | 37 | 8 |
| 71 | GCA_025190445.1.faa | 1420 | 1713 | 24 | 6 |
| 72 | GCA_025190465.1.faa | 1420 | 1648 | 30 | 14 |
| 73 | GCA_025398935.1.faa | 1420 | 1513 | 3 | 7 |
| 74 | GCA_026222675.1_.faa | 1420 | 1603 | 1 | 0 |
| 75 | GCA_028464285.1.faa | 1420 | 1526 | 21 | 3 |
| 76 | GCA_029228745.1.faa | 1420 | 1722 | 7 | 0 |
| 77 | GCA_029542285.1.faa | 1420 | 1509 | 24 | 3 |
| 78 | GCA_030480485.1.faa | 1420 | 1681 | 68 | 0 |
| 79 | GCA_030489685.1.faa | 1420 | 1634 | 43 | 0 |
| 80 | GCA_030578355.1.faa | 1420 | 1575 | 50 | 4 |
| 81 | GCA_032190575.1.faa | 1420 | 1705 | 0 | 2 |
| 82 | GCA_032190655.1.faa | 1420 | 1741 | 3 | 2 |
| 83 | GCA_032190675.1.faa | 1420 | 1718 | 3 | 0 |
| 84 | GCA_032190695.1.faa | 1420 | 1577 | 4 | 0 |
| 85 | GCA_032190735.1.faa | 1420 | 1622 | 8 | 3 |
| 86 | GCA_032195525.1.faa | 1420 | 1680 | 0 | 0 |
| 87 | GCF_001188985.1.faa | 1420 | 1675 | 4 | 2 |
| 88 | GCF_009812415.1.faa | 1420 | 1789 | 23 | 2 |
| 89 | GCF_016804305.1.faa | 1420 | 1652 | 1 | 0 |
| 90 | GCF_022701335.1.faa | 1420 | 1667 | 4 | 0 |
| 91 | GCF_027675785.1.faa | 1420 | 1652 | 0 | 0 |
| 92 | GCF_027675885.1.faa | 1420 | 1653 | 0 | 0 |
| 93 | GCF_027676485.1.faa | 1420 | 1655 | 0 | 0 |
| 94 | GCF_029813085.1.faa | 1420 | 1632 | 15 | 2 |
| 95 | GCF_030549395.1.faa | 1420 | 1618 | 11 | 1 |
| 96 | GCF_900092635.1.faa | 1420 | 1737 | 7 | 0 |

**Suppl. Table 4**: BPGA analysis results of 96 genomes of *L. pentosus* strains worldwide, which indicates a number of core genes, accessory genes, unique genes, and exclusively absent genes. The highlighted strain is the isolated strain used for the comparative study.
