## Supplementary table 5 for "Bacteriocin Diversity and Antiviral Potential of *Lactiplantibacillus pentosus from* Fermented Rice"

**Suppl. Table 5: Represents bacteriocins present in 96 genomes of L. pentosus isolated from various geographical areas.**

| **Genome No.** | **NCBI Accession**  **No.** | **Bacteriocin clusters in the genome** | | | | | **Additional bacteriocin in the same cluster** |
| --- | --- | --- | --- | --- | --- | --- | --- |
|  |  | **Pediocin** | **Plantaricin** | **Bovicin/ Putative bacteriocin** | **Pentocin** | **Sactipeptides / LAPs** |  |
| 1 | GCA_000271445.1 | - | - | - | Pentocin | - | - |
| 2 | GCA_001433755.1 | Pediocin | A | - | - | - | - |
| 3 | GCA_002211885.1 | Pediocin |  | - | - | - | - |
| 4 | GCA_002751855.1 | Pediocin | A | - | - | Sactipeptides |  |
| 5 | GCA_002850015.1 | Pediocin | NC8-alpha-beta | - | - | - | Plantaricin_A |
| 6 | GCA_002993385.1 | Pediocin |  | - | - | Sactipeptides | - |
| 7 | GCA_002993395.1 | Pediocin | NC8-alpha-beta | - | - | Sactipeptides, LAPs | Plantaricin_A |
| 8 | GCA_002993425.1 | Pediocin | A, S-alpha-beta | Bovicin | - | LAPs | - |
| 9 | GCA_002993435.1 | Pediocin | A, S-alpha-beta | Bovicin | - | Sactipeptides | - |
| 10 | GCA_002993465.1 | Pediocin | - | - | - | LAPs | - |
| 11 | GCA_002993485.1 | Pediocin | A, S-alpha-beta | Bovicin | - | - | - |
| 12 | GCA_003627295.1 | Pediocin | - | - | - | - | - |
| 13 | GCA_003627375.1 | Pediocin | - | - | - | - | - |
| 14 | GCA_003641185.1 | Pediocin | A | - | - | - | - |
| 15 | GCA_003702565.1 | Pediocin | EF | - | - | Sactipeptides | - |
| 16 | GCA_003702605.1 | Pediocin | EF | - | - | Sactipeptides | - |
| 17 | GCA_003702625.1 | Pediocin | EF | - | - | Sactipeptides | - |
| 18 | GCA_003702635.1 | Pediocin | - | Bovicin | - | Sactipeptides | - |
| 19 | GCA_003702665.1 | Pediocin | EF | - | - | Sactipeptides | - |
| 20 | GCA_004354685.1 | Pediocin | A | - | - | Sactipeptides | - |
| 21 | GCA_007991835.1 | Pediocin | A | - | - | Sactipeptides | - |
| 22 | GCA_009295675.1 | Pediocin | EF | Bovicin | - | - | - |
| 23 | GCA_017742275.1 | Pediocin | - | - | - | Sactipeptides | - |
| 24 | GCA_018069575.1 | Pediocin | - | - | Pentocin | Sactipeptides | Pentocin |
| 25 | GCA_018403455.1 | Pediocin | NC8-alpha-beta | - | - | Sactipeptides | Plantaricin_A |
| 26 | GCA_018982875.1 | Pediocin | A | - | - | Sactipeptides | - |
| 27 | GCA_018991285.1 | Pediocin | NC8-alpha-beta | - | - | Sactipeptides | Plantaricin_A |
| 28 | GCA_018991345.1 | Pediocin | NC8-alpha-beta | - | - | Sactipeptides | Plantaricin_A |
| 29 | GCA_018991355.1 | Pediocin | NC8-alpha-beta | - | - | Sactipeptides | Plantaricin_A |
| 30 | GCA_018991365.1 | Pediocin | NC8-alpha-beta | - | - | Sactipeptides | Plantaricin_A |
| 31 | GCA_018991375.1 | Pediocin | - | - | - | - | - |
| 32 | GCA_018991425.1 | Pediocin | - | - | Pentocin | Sactipeptides | - |
| 33 | GCA_018991465.1 | Pediocin | - | - | - | Sactipeptides | Pediocin |
| 34 | GCA_018991535.1 | Pediocin | A | - | - | - | - |
| 35 | GCA_018991565.1 | Pediocin | NC8-alpha-beta | Putative bacteriocin | - | Sactipeptides | Plantaricin_A |
| 36 | GCA_018993275.1 | Pediocin |  | - | Pentocin | Sactipeptides | Pentocin |
| 37 | GCA_018993345.1 | Pediocin | NC8-alpha-beta | - | - | Sactipeptides | Plantaricin_A |
| 38 | GCA_018993445.1 | Pediocin | NC8-alpha-beta | - | - | Sactipeptides | Plantaricin_A |
| 39 | GCA_018993485.1 | Pediocin | NC8-alpha-beta | - | - | Sactipeptides | Plantaricin_A |
| 40 | GCA_018993495.1 | Pediocin | NC8-alpha-beta | - | - | Sactipeptides | Plantaricin_A |
| 41 | GCA_018993525.1 | Pediocin | - |  | Pentocin | Sactipeptides | Pentocin |
| 42 | GCA_018993585.1 | Pediocin | - | - | - | Sactipeptides | - |
| 43 | GCA_018993625.1 | Pediocin | - | - | Pentocin | Sactipeptides | - |
| 44 | GCA_018993635.1 | Pediocin | NC8-alpha-beta | Putative bacteriocin | - | Sactipeptides | Plantaricin_A |
| 45 | GCA_018993665.1 | Pediocin |  | - | - | Sactipeptides |  |
| 46 | GCA_018993725.1 | Pediocin | NC8-alpha-beta | - | - | Sactipeptides | Plantaricin_A |
| 47 | GCA_018993945.1 | Pediocin | NC8-alpha-beta | - | - | Sactipeptides | Plantaricin_A |
| 48 | GCA_018993965.2 | Pediocin |  | - | - | - | - |
| 49 | GCA_020181715.1 | - | NC8-alpha-beta | - | - | Sactipeptides | Plantaricin_A |
| 50 | GCA_020532005.1 | Pediocin |  | - | - | Sactipeptides |  |
| 51 | GCA_021384425.1 | Pediocin | NC8-alpha-beta | - | - | Sactipeptides | Plantaricin_A |
| 52 | GCA_022936785.1 | - | NC8-alpha-beta | - | - | Sactipeptides | Plantaricin_A |
| 53 | GCA_022936805.1 | Pediocin |  | Bovicin | - | - | - |
| 54 | GCA_022936825.1 | Pediocin | A |  |  | Sactipeptides |  |
| 55 | GCA_023823145.1 | Pediocin |  | - | - | - | - |
| 56 | GCA_023972895.1 | Pediocin | A | - | - | - | - |
| 57 | GCA_023980805.1 | Pediocin | A | - | - | - | - |
| 58 | GCA_025122375.1 | Pediocin | - | - | - | Sactipeptides | - |
| 59 | GCA_025129205.1 | 2 Pediocin | - | - | - | Sactipeptides | - |
| 60 | GCA_025188445.1 | Pediocin | A,  NC8-alpha-beta | - | - | Sactipeptides | - |
| 61 | GCA_025190245.1 | Pediocin | NC8-alpha-beta | - | - | Sactipeptides | Plantaricin_A |
| 62 | GCA_025190265.1 | Pediocin | A |  |  | Sactipeptides | - |
| 63 | GCA_025190285.1 | Pediocin | - | - | - | Sactipeptides | - |
| 64 | GCA_025190295.1 | - | A | - | - | Sactipeptides | - |
| 65 | GCA_025190325.1 | Pediocin | A | - | - | Sactipeptides | - |
| 66 | GCA_025190345.1 | Pediocin | A | Bovicin | - | Sactipeptides | - |
| 67 | GCA_025190365.1 | Pediocin | A | - | - | Sactipeptides | - |
| 68 | GCA_025190375.1 | Pediocin | EF | - | - | Sactipeptides | - |
| 69 | GCA_025190395.1 | Pediocin | A, S-alpha-beta | Bovicin | - | Sactipeptides | - |
| 70 | GCA_025190425.1 | Pediocin | - | - | - | Sactipeptides | Pediocin |
| 71 | GCA_025190445.1 | Pediocin | A | - | - | Sactipeptides | - |
| 72 | GCA_025190465.1 | Pediocin | - | - | - | Sactipeptides | - |
| 73 | GCA_025398935.1 | Pediocin | EF | - | - | - | - |
| 74 | GCA_026222675.1 | Pediocin | A |  |  |  |  |
| 75 | GCA_028464285.1 | Pediocin | - | - | - | - | - |
| 76 | GCA_029228745.1 | Pediocin | A | - | - | LAPs | - |
| 77 | GCA_029542285.1 | Pediocin | - | - | - | - | - |
| 78 | GCA_030480485.1 | Pediocin | A | - | - | - | - |
| 79 | GCA_030489685.1 | Pediocin | - | - | - | - | - |
| 80 | GCA_030578355.1 | 2 Pediocin | EF | - | - | Sactipeptides | - |
| 81 | GCA_032190575.1 | Pediocin | - | - | Pentocin | - | Pentocin |
| 82 | GCA_032190655.1 | Pediocin | - | - | Pentocin | - | Pentocin |
| 83 | GCA_032190675.1 | Pediocin | - | - | - | - | Pediocin |
| 84 | GCA_032190695.1 | Pediocin | NC8-alpha-beta | - | - | - | Plantaricin_A |
| 85 | GCA_032190735.1 | Pediocin | NC8-alpha-beta | Putative bacteriocin | - | - | Plantaricin_A |
| 86 | GCA_032195525.1 | Pediocin | A | - | - | - | - |
| 87 | GCF_001188985.1 | Pediocin | A | - |  | Sactipeptides |  |
| 88 | GCF_009812415.1 | Pediocin | - | - | Pentocin | - | Acidocin |
| 89 | GCF_016804305.1 | Pediocin | A | - | - | - | - |
| 90 | GCF_022701335.1 | Pediocin | A | - | - | - | - |
| 91 | GCF_027675785.1 | Pediocin | A | Bovicin |  | - | - |
| 92 | GCF_027675885.1 | Pediocin | A | Bovicin |  | Sactipeptides | - |
| 93 | GCF_027676485.1 | Pediocin | A | Bovicin | - | - | - |
| 94 | GCF_029813085.1 | - | EF | - | - | Sactipeptides | - |
| 95 | GCF_030549395.1 | 2 Pediocin | - | - | - | Sactipeptides | - |
| 96 | GCF_900092635.1 | Pediocin | A | - | - | - | - |
