## Supplementary table 6 for "Bacteriocin Diversity and Antiviral Potential of *Lactiplantibacillus pentosus from* Fermented Rice"

**Suppl. Table 6:** Represents the C-score, TM-score, and corresponding PDB hits for the predicted structures of three bacteriocins—Pentoplantaricin_EF, Pentopediocin, and Pentobovicin—as generated by I-TASSER.

| **Sl. No.** | **Ligand** | **C-score** | **TM-score** | **PDB Hit** |
| --- | --- | --- | --- | --- |
| 1. | Pentoplantaricin_EF | -2.37 | 0.44±0.14 | 2RLW |
| 2. | Pentopediocin | 0.59 | 0.64±0.14 | 2IP6 |
| 3. | Pentobovicin, | -2.64 | 0.41±0.14 | 5LFI |
