## Supplementary table 7 for "Bacteriocin Diversity and Antiviral Potential of *Lactiplantibacillus pentosus from* Fermented Rice"

| **Sl. No.** | **Ligand** | **Target** | **Target PDB ID** | **Binding energy (kcal/mol)** |
| --- | --- | --- | --- | --- |
| 1. | Pentoplantaricin_EF | Prefusion 2019-nCoV spike glycoprotein | 6VSB | -1439.4 |
| 2. | Pentoplantaricin_EF | SARS-CoV-2 Omicron spike protein | 7T9J | -1114.9 |
| 3. | Pentoplantaricin_EF | Hepatitis E Virus Capsid Protein E2s Domain | 3RKC | -881.3 |
| 4. | Pentopediocin | Prefusion 2019-nCoV spike glycoprotein | 6VSB | -887.8 |
| 5. | Pentopediocin | SARS-CoV-2 Omicron spike protein | 7T9J | -975.2 |
| 6. | Pentopediocin | Hepatitis E Virus Capsid Protein E2s Domain | 3RKC | -869.2 |
| 7. | Pentobovicin | Prefusion 2019-nCoV spike glycoprotein | 6VSB | -971.7 |
| 8. | Pentobovicin | SARS-CoV-2 Omicron spike protein | 7T9J | -1153.4 |
| 9. | Pentobovicin | Hepatitis E Virus Capsid Protein E2s Domain | 3RKC | -928.8 |

**Suppl. Table 7:** Summary of binding interactions between bacteriocins and viral targets and their corresponding binding energies for each target-ligand pair.
