## Supplementary material for "Bacteriocin Diversity and Antiviral Potential of *Lactiplantibacillus pentosus from* Fermented Rice": statistical data

| | | Group A | Group B | Group C | Group D | Group E | Group F | Group G | Group H | Group I |
|---|---|---|---|---|---|---|---|---|---|---|
| | | MRS | BHI | NB | LB | YPD | M17 | SB | SD | ELK |
| 1 | Title | 0.21650 | 0.23750 | 0.099 | 0.084 | 0.16550 | 0.500 | 0.3275 | 0.0830 | 0.13150 |
| 2 | Title | 0.19854 | 0.21210 | 0.080 | 0.097 | 0.26558 | 0.485 | 0.3125 | 0.0940 | 0.21000 |
| 3 | Title | 0.25400 | 0.28500 | 0.070 | 0.078 | 0.27890 | 0.458 | 0.4150 | 0.0725 | 0.12000 |

| Table format: **Column** | | Group A | Group B | Group C | Group D | Group E | Group F | Group G | Group H |
|---|---|---|---|---|---|---|---|---|---|
| | | MRS | BHI | NB | LB | YPD | M17 | SB | SD |
| 1 | Title | 13.0 | 0 | 0 | 0 | 11 | 12 | 0 | 0 |
| 2 | Title | 12.0 | 0 | 0 | 0 | 11 | 11 | 0 | 0 |
| 3 | Title | 13.0 | 0 | 0 | 0 | 11 | 12 | 0 | 0 |

| | Group I | Group J | Group K |
|---|---|---|---|
| | ELK | Streptomycin | Nisin |
| 1 | 10.0 | 16 | 0 |
| 2 | 11.0 | 16 | 0 |
| 3 | 11.0 | 15 | 0 |

| Table format: **Column** | | Group A | Group B | Group C | Group D | Group E | Group F | Group G | Group H | Group I |
|---|---|---|---|---|---|---|---|---|---|---|
| | | -20 | 4 | 30 | 40 | 50 | 60 | 70 | 80 | 90 |
| 1 | Title | 15 | 15.0 | 14 | 14 | 13.0 | 14 | 15 | 15 | 15 |
| 2 | Title | 15 | 14.0 | 14 | 12 | 13.0 | 14 | 15 | 15 | 15 |
| 3 | Title | 15 | 15.0 | 14 | 14 | 12.0 | 13 | 14 | 15 | 15 |

| | Group J | Group K | Group L |
|---|---|---|---|
| | 100 | 121 | NMRS |
| 1 | 15 | 16 | 13 |
| 2 | 15 | 16 | 13 |
| 3 | 14 | 16 | 13 |

| Table format: **Column** | | Group A | Group B | Group C | Group D | Group E | Group F | Group G | Group H |
|---|---|---|---|---|---|---|---|---|---|
| | | 3 | 4 | 5 | 6 | 7 | 8 | 9 | 10 |
| 1 | Title | 12 | 11.0 | 0 | 0 | 0 | 0 | 0 | 0 |
| 2 | Title | 12 | 10.0 | 0 | 0 | 0 | 0 | 0 | 0 |
| 3 | Title | 12 | 11.0 | 0 | 0 | 0 | 0 | 0 | 0 |

| | Group I | Group J |
|---|---|---|
| | 11 | N |
| 1 | 0 | 12 |
| 2 | 0 | 11 |
| 3 | 0 | 12 |

| Table format: **Column** | | Group A | Group B | Group C | Group D | Group E | Group F | Group G | Group H | Group I |
|---|---|---|---|---|---|---|---|---|---|---|
| | | $FeSO_4$ | $CuCl_2$ | $CaCl_2$ | KCl | $ZnSO_4$ | $MgSO_4$ | $MnSO_4$ | N-MRS | Streptomycin |
| 1 | Title | 13 | 12 | 13 | 14 | 14 | 13 | 14 | 13 | 17 |
| 2 | Title | 13 | 12 | 13 | 14 | 14 | 14 | 14 | 14 | 16 |
| 3 | Title | 12 | 12 | 13 | 14 | 13 | 13 | 14 | 14 | 17 |

| | Group J |
|---|---|
| | Nisin |
| 1 | 0 |
| 2 | 0 |
| 3 | 0 |

| Table format: **Column** | | Group A | Group B | Group C | Group D | Group E | Group F | Group G | Group H | Group I |
|---|---|---|---|---|---|---|---|---|---|---|
| | | Protease | Proteinase K | Trypsin | Chymotrypsin | Lysozyme | Amylase | Lipase | Catalase | Pepsin |
| 1 | Title | 13 | 12 | 11 | 13 | 12 | 13 | 13 | 13.0 | 13 |
| 2 | Title | 13 | 12 | 11 | 13 | 12 | 13 | 12 | 11.0 | 12 |
| 3 | Title | 12 | 11 | 12 | 13 | 12 | 12 | 13 | 10.0 | 13 |

| | Group J | Group K | Group L |
|---|---|---|---|
| | N-MRS | Streptomycin | Nisin |
| 1 | 13 | 16 | 0 |
| 2 | 13 | 15 | 0 |
| 3 | 12 | 16 | 0 |

| Table format: **Column** | | Group A | Group B | Group C | Group D | Group E | Group F | Group G | Group H | Group I |
|---|---|---|---|---|---|---|---|---|---|---|
| | | Acetone | Chloroform | Ethanol | Methanol | DMSO | Urea | EDTA | SDS | TCA |
| 1 | Title | 13 | 14 | 15 | 14 | 16 | 16 | 18 | 14 | 14 |
| 2 | Title | 13 | 14 | 15 | 14 | 16 | 15 | 18 | 13 | 12 |
| 3 | Title | 13 | 14 | 15 | 14 | 15 | 14 | 17 | 14 | 14 |

| | Group J | Group K | Group L | Group M | Group N | Group O |
|---|---|---|---|---|---|---|
| | Tween-20 | Tween-80 | Triton -X-100 | N-MRS | Streptomycin | Nisin |
| 1 | 22 | 16 | 17 | 18 | 17 | 0 |
| 2 | 21 | 17 | 17 | 18 | 17 | 0 |
| 3 | 23 | 14 | 17 | 18 | 16 | 0 |

| | | X | Group A | | Group B | |
|---|---|---|---|---|---|---|
| | | Time (hour) | OD at 600 nm | | Zone of Inhibition (mm) | |
| | | X | A:Y1 | A:Y2 | B:Y1 | B:Y2 |
| 1 | Title | 0.0 | 0.005 | 0.007 | 0.00 | 0.00 |
| 2 | Title | 1.0 | 0.007 | 0.008 | 0.00 | 0.00 |
| 3 | Title | 2.0 | 0.010 | 0.009 | 0.00 | 0.00 |
| 4 | Title | 3.0 | 0.014 | 0.012 | 0.00 | 0.00 |
| 5 | Title | 4.0 | 0.019 | 0.017 | 0.00 | 0.00 |
| 6 | Title | 5.0 | 0.024 | 0.024 | 0.00 | 0.00 |
| 7 | Title | 6.0 | 0.046 | 0.042 | 0.00 | 0.00 |
| 8 | Title | 7.0 | 0.056 | 0.050 | 0.00 | 0.00 |
| 9 | Title | 8.0 | 0.076 | 0.069 | 0.00 | 0.00 |
| 10 | Title | 9.0 | 0.153 | 0.121 | 0.00 | 0.00 |
| 11 | Title | 10.0 | 0.220 | 0.215 | 0.00 | 0.00 |
| 12 | Title | 11.0 | 0.410 | 0.560 | 0.00 | 0.00 |
| 13 | Title | 12.0 | 0.730 | 0.730 | 0.00 | 0.00 |
| 14 | Title | 13.0 | 0.780 | 0.970 | 10.00 | 10.00 |
| 15 | Title | 14.0 | 0.855 | 1.210 | 10.00 | 10.00 |
| 16 | Title | 15.0 | 1.180 | 1.400 | 11.00 | 11.00 |
| 17 | Title | 16.0 | 1.320 | 1.430 | 11.00 | 11.00 |
| 18 | Title | 17.0 | 1.550 | 1.640 | 12.00 | 12.00 |
| 19 | Title | 18.0 | 1.811 | 1.720 | 12.00 | 12.00 |
| 20 | Title | 19.0 | 1.813 | 1.820 | 13.00 | 14.00 |
| 21 | Title | 20.0 | 2.020 | 1.960 | 13.00 | 14.00 |
| 22 | Title | 21.0 | 1.700 | 1.810 | 14.00 | 14.00 |
| 23 | Title | 22.0 | 2.350 | 2.080 | 15.00 | 15.00 |
| 24 | Title | 23.0 | 2.360 | 2.290 | 15.00 | 15.00 |
| 25 | Title | 24.0 | 2.440 | 2.360 | 16.00 | 16.00 |
| 26 | Title | 25.0 | 3.010 | 3.340 | 16.00 | 16.00 |
| 27 | Title | 26.0 | 3.360 | 3.280 | 16.00 | 17.00 |
| 28 | Title | 27.0 | 3.510 | 3.180 | 16.00 | 17.00 |
| 29 | Title | 28.0 | 3.140 | 3.140 | 17.00 | 18.00 |
| 30 | Title | 29.0 | 3.120 | 3.110 | 18.00 | 19.00 |
| 31 | Title | 30.0 | 2.940 | 2.830 | 19.00 | 19.00 |
| 32 | Title | 31.0 | 2.360 | 2.820 | 19.00 | 19.00 |
| 33 | Title | 32.0 | 2.540 | 2.760 | 19.00 | 19.00 |
| 34 | Title | 33.0 | 2.560 | 2.680 | 19.00 | 19.00 |
| 35 | Title | 34.0 | 2.695 | 2.710 | 19.00 | 19.00 |
| 36 | Title | 35.0 | | | | |
| 37 | Title | 36.0 | | | | |

| 2way ANOVA | | | | | | |
|---|---|---|---|---|---|---|
| 1 | Table Analyzed | Data 1 | | | | |
| 2 | | | | | | |
| 3 | **Two-way ANOVA** | Ordinary | | | | |
| 4 | Alpha | 0.05 | | | | |
| 5 | | | | | | |
| 6 | **Source of Variation** | **% of total variation** | **P value** | **P value summary** | **Significant?** | |
| 7 | Row Factor | 0.4240 | 0.5160 | ns | No | |
| 8 | Column Factor | 94.66 | <0.0001 | **** | Yes | |
| 9 | | | | | | |
| 10 | **ANOVA table** | **SS** | **DF** | **MS** | **F (DFn, DFd)** | **P value** |
| 11 | Row Factor | 0.001961 | 2 | 0.0009807 | F (2, 16) = 0.6897 | P=0.5160 |
| 12 | Column Factor | 0.4379 | 8 | 0.05473 | F (8, 16) = 38.49 | P<0.0001 |
| 13 | Residual | 0.02275 | 16 | 0.001422 | | |

| 2way ANOVA | | | | | | |
|---|---|---|---|---|---|---|
| 1 | Table Analyzed | Data 2 | | | | |
| 2 | | | | | | |
| 3 | **Two-way ANOVA** | Ordinary | | | | |
| 4 | Alpha | 0.05 | | | | |
| 5 | | | | | | |
| 6 | **Source of Variation** | **% of total variation** | **P value** | **P value summary** | **Significant?** | |
| 7 | Row Factor | 0.004677 | 0.7946 | ns | No | |
| 8 | Column Factor | 99.79 | <0.0001 | **** | Yes | |
| 9 | | | | | | |
| 10 | **ANOVA table** | **SS** | **DF** | **MS** | **F (DFn, DFd)** | **P value** |
| 11 | Row Factor | 0.06061 | 2 | 0.03030 | F (2, 20) = 0.2326 | P=0.7946 |
| 12 | Column Factor | 1293 | 10 | 129.3 | F (10, 20) = 992.5 | P<0.0001 |
| 13 | Residual | 2.606 | 20 | 0.1303 | | |

**2way ANOVA**

| | | | | | | |
|---|---|---|---|---|---|---|
| 1 | Table Analyzed | Data 3 | | | | |
| 2 | | | | | | |
| 3 | **Two-way ANOVA** | Ordinary | | | | |
| 4 | Alpha | 0.05 | | | | |
| 5 | | | | | | |
| 6 | **Source of Variation** | **% of total variation** | **P value** | **P value summary** | **Significant?** | |
| 7 | Row Factor | 1.919 | 0.2439 | ns | No | |
| 8 | Column Factor | 84.06 | <0.0001 | **** | Yes | |
| 9 | | | | | | |
| 10 | **ANOVA table** | **SS** | **DF** | **MS** | **F (DFn, DFd)** | **P value** |
| 11 | Row Factor | 0.7222 | 2 | 0.3611 | F (2, 22) = 1.505 | P=0.2439 |
| 12 | Column Factor | 31.64 | 11 | 2.876 | F (11, 22) = 11.99 | P<0.0001 |
| 13 | Residual | 5.278 | 22 | 0.2399 | | |

| 2way ANOVA | | | | | | |
|---|---|---|---|---|---|---|
| 1 | Table Analyzed | Data 4 | | | | |
| 2 | | | | | | |
| 3 | **Two-way ANOVA** | Ordinary | | | | |
| 4 | Alpha | 0.05 | | | | |
| 5 | | | | | | |
| 6 | **Source of Variation** | **% of total variation** | **P value** | **P value summary** | **Significant?** | |
| 7 | Row Factor | 0.03215 | 0.1342 | ns | No | |
| 8 | Column Factor | 99.84 | <0.0001 | **** | Yes | |
| 9 | | | | | | |
| 10 | **ANOVA table** | **SS** | **DF** | **MS** | **F (DFn, DFd)** | **P value** |
| 11 | Row Factor | 0.2667 | 2 | 0.1333 | F (2, 18) = 2.250 | P=0.1342 |
| 12 | Column Factor | 828.0 | 9 | 92.00 | F (9, 18) = 1553 | P<0.0001 |
| 13 | Residual | 1.067 | 18 | 0.05926 | | |

| 2way ANOVA | | | | | | |
|---|---|---|---|---|---|---|
| 1 | Table Analyzed | Data 5 | | | | |
| 2 | | | | | | |
| 3 | **Two-way ANOVA** | Ordinary | | | | |
| 4 | Alpha | 0.05 | | | | |
| 5 | | | | | | |
| 6 | **Source of Variation** | **% of total variation** | **P value** | **P value summary** | **Significant?** | |
| 7 | Row Factor | 0.03648 | 0.5730 | ns | No | |
| 8 | Column Factor | 99.39 | <0.0001 | **** | Yes | |
| 9 | | | | | | |
| 10 | **ANOVA table** | **SS** | **DF** | **MS** | **F (DFn, DFd)** | **P value** |
| 11 | Row Factor | 0.2000 | 2 | 0.1000 | F (2, 18) = 0.5745 | P=0.5730 |
| 12 | Column Factor | 545.0 | 9 | 60.55 | F (9, 18) = 347.9 | P<0.0001 |
| 13 | Residual | 3.133 | 18 | 0.1741 | | |

| 2way ANOVA | | | | | | |
|---|---|---|---|---|---|---|
| 1 | Table Analyzed | Data 6 | | | | |
| 2 | | | | | | |
| 3 | **Two-way ANOVA** | Ordinary | | | | |
| 4 | Alpha | 0.05 | | | | |
| 5 | | | | | | |
| 6 | **Source of Variation** | **% of total variation** | **P value** | **P value summary** | **Significant?** | |
| 7 | Row Factor | 0.3537 | 0.1250 | ns | No | |
| 8 | Column Factor | 97.95 | <0.0001 | **** | Yes | |
| 9 | | | | | | |
| 10 | **ANOVA table** | **SS** | **DF** | **MS** | **F (DFn, DFd)** | **P value** |
| 11 | Row Factor | 1.722 | 2 | 0.8611 | F (2, 22) = 2.289 | P=0.1250 |
| 12 | Column Factor | 477.0 | 11 | 43.36 | F (11, 22) = 115.2 | P<0.0001 |
| 13 | Residual | 8.278 | 22 | 0.3763 | | |

| 2way ANOVA | | | | | | |
|---|---|---|---|---|---|---|
| 1 | Table Analyzed | Data 7 | | | | |
| 2 | | | | | | |
| 3 | **Two-way ANOVA** | Ordinary | | | | |
| 4 | Alpha | 0.05 | | | | |
| 5 | | | | | | |
| 6 | **Source of Variation** | **% of total variation** | **P value** | **P value summary** | **Significant?** | |
| 7 | Row Factor | 0.1331 | 0.2716 | ns | No | |
| 8 | Column Factor | 98.50 | <0.0001 | **** | Yes | |
| 9 | | | | | | |
| 10 | **ANOVA table** | **SS** | **DF** | **MS** | **F (DFn, DFd)** | **P value** |
| 11 | Row Factor | 1.244 | 2 | 0.6222 | F (2, 28) = 1.366 | P=0.2716 |
| 12 | Column Factor | 921.2 | 14 | 65.80 | F (14, 28) = 144.4 | P<0.0001 |
| 13 | Residual | 12.76 | 28 | 0.4556 | | |

| 2way ANOVA | | | | | |
|---|---|---|---|---|---|
| 1 | Table Analyzed | Exponential decay | | | |
| 2 | | | | | |
| 3 | **Two-way ANOVA** | Ordinary | | | |
| 4 | Alpha | 0.05 | | | |
| 5 | | | | | |
| 6 | **Source of Variation** | **% of total variation** | **P value** | **P value summary** | **Significant?** |
| 7 | Interaction | 23.25 | <0.0001 | **** | Yes |
| 8 | Row Factor | 42.08 | <0.0001 | **** | Yes |
| 9 | Column Factor | 34.62 | <0.0001 | **** | Yes |
| 10 | | | | | |
| 11 | **ANOVA table** | **SS** | **DF** | **MS** | **F (DFn, DFd)** |
| 12 | Interaction | 1517 | 34 | 44.63 | F (34, 70) = 909.1 |
| 13 | Row Factor | 2746 | 34 | 80.78 | F (34, 70) = 1645 |
| 14 | Column Factor | 2259 | 1 | 2259 | F (1, 70) = 46017 |
| 15 | Residual | 3.437 | 70 | 0.04910 | |
| 16 | | | | | |
| 17 | **Difference between column means** | | | | |
| 18 | Mean of OD at 600 nm | 1.480 | | | |
| 19 | Mean of Zone of Inhibition (mm) | 9.514 | | | |
| 20 | Difference between means | -8.034 | | | |
| 21 | SE of difference | 0.03745 | | | |
| 22 | 95% CI of difference | -8.109 to -7.960 | | | |

| | |
|---|---|
| 1 | |
| 2 | |
| 3 | |
| 4 | |
| 5 | |
| 6 | |
| 7 | |
| 8 | |
| 9 | |
| 10 | |
| 11 | **P value** |
| 12 | P<0.0001 |
| 13 | P<0.0001 |
| 14 | P<0.0001 |
| 15 | |
| 16 | |
| 17 | |
| 18 | |
| 19 | |
| 20 | |
| 21 | |
| 22 | |

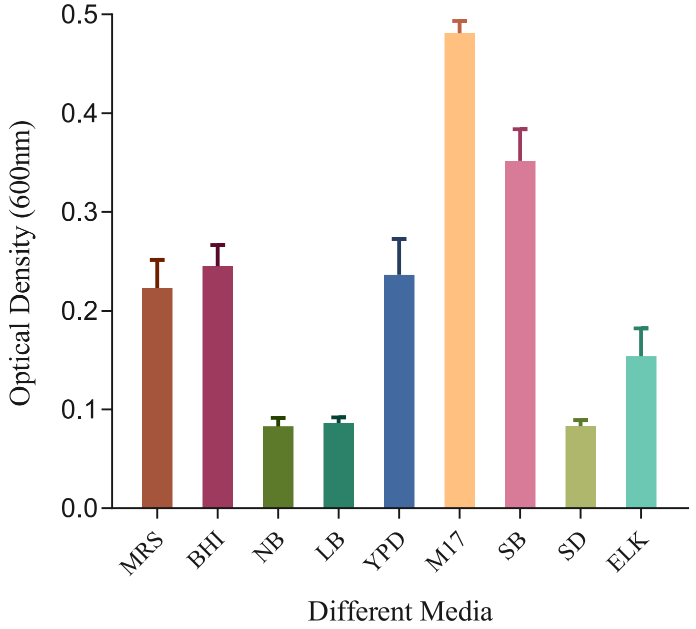

0.5
0.4
0.3
0.2
0.1
0.0
Optical Density (600nm)
MRS
BHI
NB
LB
YPD
M17
SB
SD
ELK
Different Media

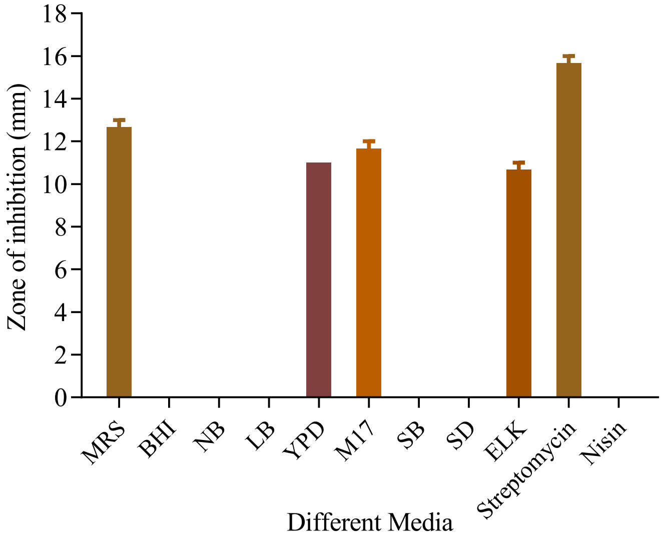

18
16
14
12
10
8
6
4
2
0
Zone of inhibition (mm)
MRS
BHI
NB
LB
YPD
M17
SB
SD
ELK
Streptomycin
Nisin
Different Media

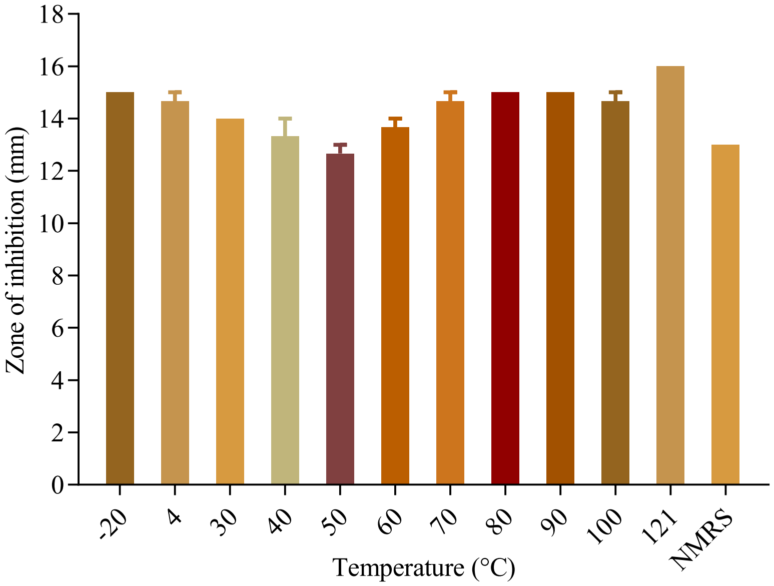

Zone of inhibition (mm)
0
2
4
6
8
10
12
14
16
18
-20
4
30
40
50
60
70
80
90
100
121
NMRS
Temperature (°C)

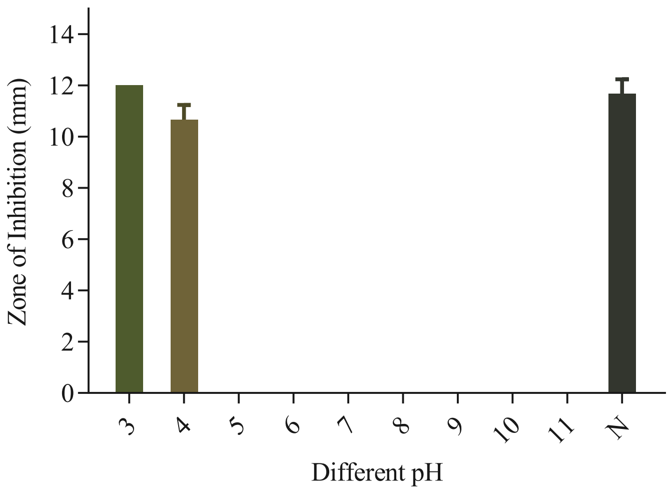
Zone of Inhibition (mm)
14
12
10
8
6
4
2
0
3
4
5
6
7
8
9
10
11
N
Different pH

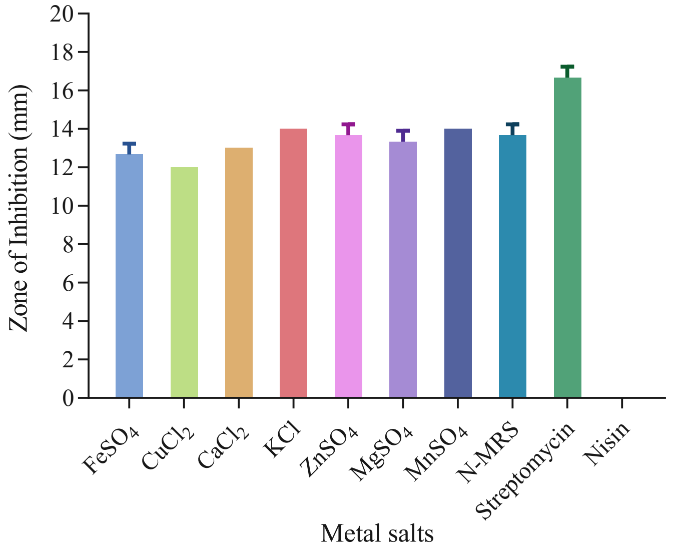

20
18
16
14
12
10
8
6
4
2
0
Zone of Inhibition (mm)
FeSO4
CuCl2
CaCl2
KCl
ZnSO4
MgSO4
MnSO4
N-MRS
Streptomycin
Nisin
Metal salts

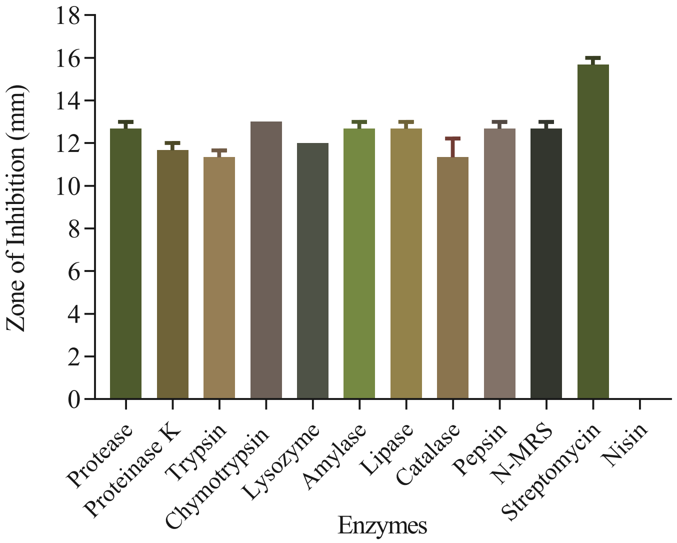
Zone of Inhibition (mm)
0
2
4
6
8
10
12
14
16
18
Protease
Proteinase K
Trypsin
Chymotrypsin
Lysozyme
Amylase
Lipase
Catalase
Pepsin
N-MRS
Streptomycin
Nisin
Enzymes

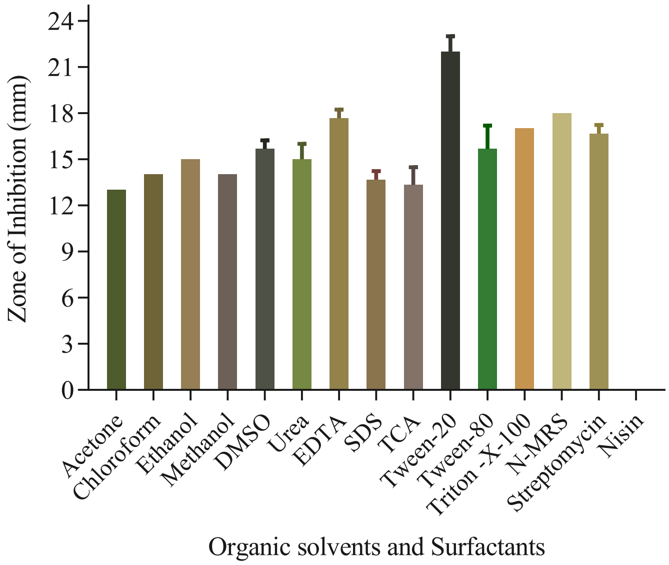

Zone of Inhibition (mm)
24
21
18
15
12
9
6
3
0
Acetone
Chloroform
Ethanol
Methanol
DMSO
Urea
EDTA
SDS
TCA
Tween-20
Tween-80
Triton -X-100
N-MRS
Streptomycin
Nisin
Organic solvents and Surfactants

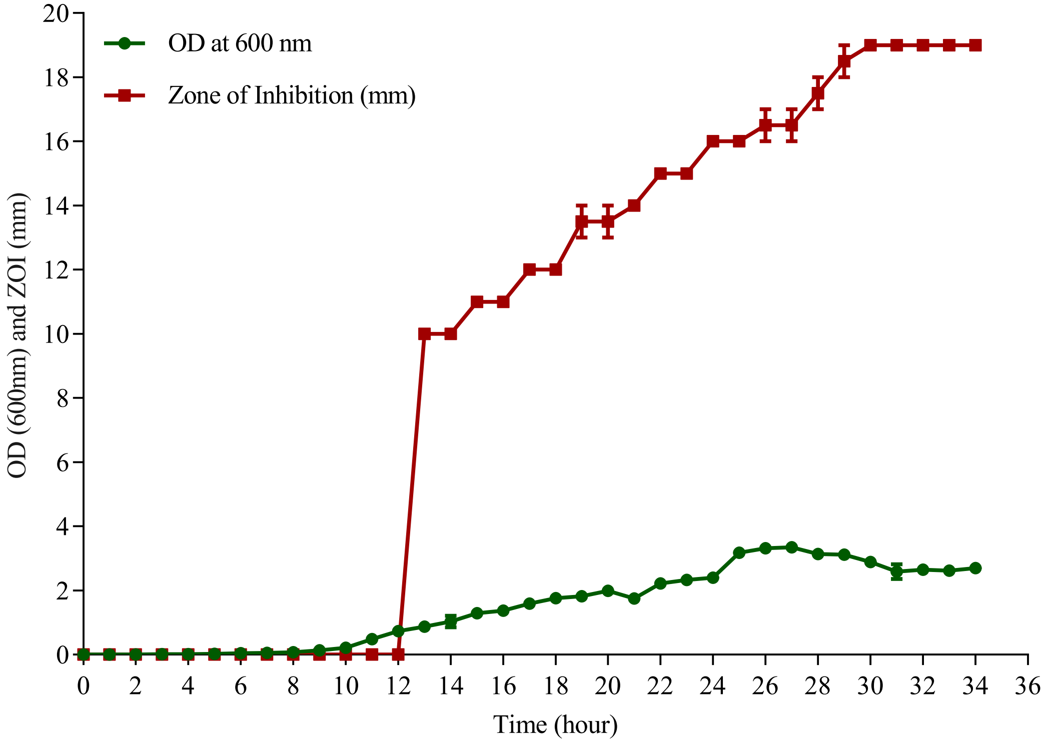

OD at 600 nm
Zone of Inhibition (mm)
OD (600nm) and ZOI (mm)
Time (hour)
0
2
4
6
8
10
12
14
16
18
20
22
24
26
28
30
32
34
36

| Constant | Value |
|---|---|
| Experiment Da | 16-Jan-23 |
| Experiment ID | |
| Notebook ID | |
| Project | |
| Experimenter | |
| Protocol | |
